# Diversity matters: Deep-sea mussels harbor multiple symbiont strains

**DOI:** 10.1101/531459

**Authors:** Rebecca Ansorge, Stefano Romano, Lizbeth Sayavedra, Anne Kupczok, Halina E. Tegetmeyer, Nicole Dubilier, Jillian Petersen

**Author notes:** These authors contributed equally (Stefano Romano, Lizbeth Sayavedra).

## Abstract

Genetic diversity of closely-related free-living microbes is widespread and underpins ecosystem functioning, but most evolutionary theories predict that it destabilizes intimate mutualisms. Indeed, symbiont strain diversity has long assumed to be restricted in intracellular bacteria associated with animals. Here, we sequenced the metagenomes and metatranscriptomes of 18 *Bathymodiolus* mussel individuals from four species, covering their known distribution range at deep-sea hydrothermal vents in the Atlantic. We show that as many as 16 strains of intracellular, sulfur-oxidizing symbionts coexist in individual *Bathymodiolus* mussels. Co-occurring symbiont strains differed extensively in key metabolic functions, such as the use of energy and nutrient sources, electron acceptors and viral defense mechanisms. Most strain-specific genes were expressed, highlighting their adaptive potential. We show that fine-scale diversity is pervasive in *Bathymodiolus* symbionts, and hypothesize that it may be widespread in low-cost symbioses where the environment, not the host, feeds the symbionts.

## Introduction

Within-species variability is ubiquitous in natural bacterial populations and occurs at many levels, from single nucleotide polymorphisms (SNPs) to differences in gene content and regulation. These fine-scale differences can have major functional consequences and thus define microbial lifestyles. For example, a single regulatory gene or a mutation can dramatically alter the host range of bacterial symbionts and human pathogens^1,2^. In the human gut microbiome, gene copy number variation among different strains of the same bacterial species is linked to host desease^3^. However, many of these functional differences are invisible at the level of marker genes commonly used in microbiome studies, such as the gene encoding 16S rRNA.

In free-living microbial communities, diversity underpins ecosystem functioning and resilience^4,5^. However, in symbiotic associations, genetic diversity of microbes within host individuals can destabilize relationships between hosts and their symbionts. This is because diversity can lead to increased conflict between hosts and symbionts, and among co-existing symbionts within single individuals^6^. These inherent evolutionary conflicts can be alleviated by stabilizing mechanisms such as vertical transmission, partner choice and sanctioning, ensuring partner fidelity, or allowing hosts to discriminate against low quality partners^7–9^. These stabilizing mechanisms are hypothesized to explain the remarkably restricted diversity of symbionts in a range of associations from aphids with their *Buchnera* endosymbionts to legume nodules that contain only a single strain of rhizobial symbiont. High-throughput sequencing of natural symbiont populations is beginning to uncover unexpected within-species diversity, despite low diversity at the species level^10–13^. But does such within-species symbiont diversity bear a cost to the host? While higher diversity may create conflicts among symbionts residing in a single host, it may also bring benefits to hosts by allowing them to access a range of functions^14,15^. However, it is not understood under which conditions within-species symbiont diversity is beneficial to hosts, and efforts to understand the evolutionary implications of complex host-associated communities are in their infancy^16,17^.

Metagenomes are essential for understanding natural within-species diversity, how such diversity evolves, and how it affects function, particularly in uncultivable organisms. However, teasing apart highly similar strain genomes in metagenomes remains a major challenge^18–20^. Deep-sea *Bathymodiolus* mussels are ideal for investigating the functional and evolutionary implications of symbiont strain diversity, as they host only two bacterial symbiont species: One sulfur-oxidizing (SOX), and one methane-oxidizing (MOX) symbiont^21–23^. These symbionts co-occur inside specialized gill epithelial cells called bacteriocytes and use reduced compounds from hydrothermal fluids as energy sources for carbon fixation. The symbionts thus provide their hosts with nutrition in the nutrient-poor deep sea, allowing these mussels to dominate hydrothermal vent and cold seep communities worldwide^21–23^.

The SOX symbionts of *Bathymodiolus* are very closely related to a ubiquitous group of free-living bacteria called SUP05, and their symbioses with deep-sea mussels have likely evolved multiple times from within the SUP05 clade^24^. With few exceptions, each *Bathymodiolus* host harbors a single 16S SOX symbiont phylotype^25,26^. However, studies of the more variable ribosomal internal transcribed spacer indicated that more than one symbiont strain may colonize individual mussels^27,28^. Metagenomics of one *Bathymodiolus* species recently showed that ‘subpopulations’ of SOX symbionts differed in key functions such as hydrogen oxidation and nitrate respiration^29^. These observations raise a number of questions: How widespread is strain diversity, how many strains co-exist in a host individual, and how is such fine-scale diversity stably maintained in symbiosis over evolutionary time^30^?To address these questions, we performed high-resolution metagenomic and metatranscriptomic analyses of the symbiont populations of 18 host individuals from four *Bathymodiolus* species that were collected from four geochemically distinct, hydrothermal vents along the Mid-Atlantic Ridge.

## Results and Discussion

### Genome-wide symbiont heterogeneity

We assembled Illumina metagenomes and used differential coverage and contig connectivity data to retrieve a consensus reference genome of the *Bathymodiolus* SOX symbiont for each vent field and host species (from each vent field only one host species was found, see Methods) (Fig. 1). The symbiont bins ranged from 2 to 3 Mbp and were ≥ 94% complete (Tab. S1). In 12 out of 18 host individuals we did not detect any SNPs in the symbiont 16S rRNA genes. In the other six, we detected low-frequency SNPs, present in 8-16% of the symbiont population and some SNPs appeared in more than one individual (Extended Data Tab. 1). This supports a previous study detecting low-abundance SOX 16S rRNA phylotypes in some host individuals that are closely related to the known *Bathymodiolus* symbionts (> 98.8% similarity)^23^.

**Figure 1.**
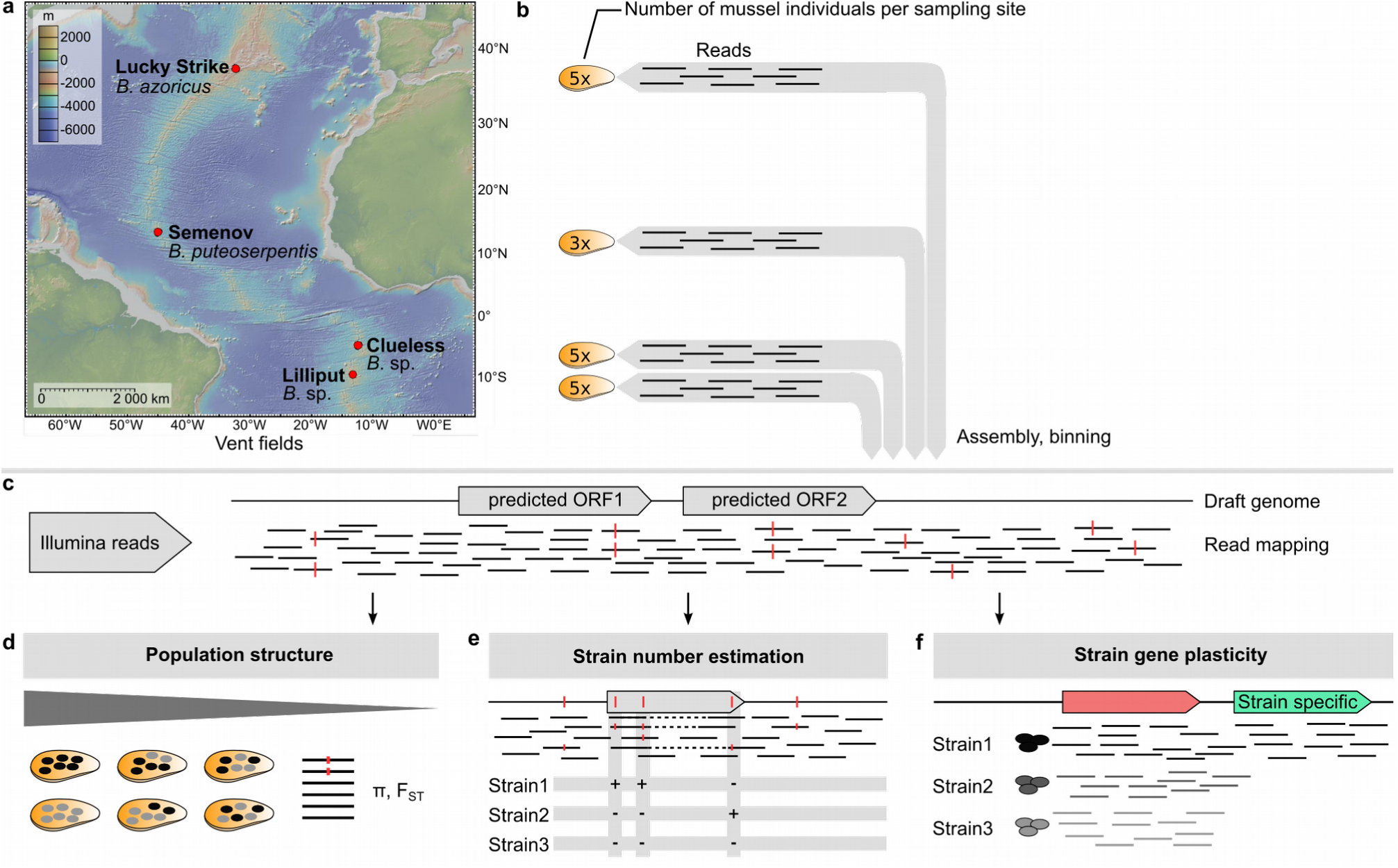
Overview of the workflow developed for this study. **(a)** *Bathymodiolus* mussels were sampled at four vent fields along the Mid-Atlantic Ridge (MAR), **(b)** metagenomes of the sulfur-oxidizing symbiont (SOX) were assembled as a consensus for each site, binned and annotated, **(c)** for each sample, reads were mapped to per-site consensus draft genomes. We analyzed three aspects of symbiont strain diversity: **(d)** symbiont population structure by single nucleotide polymorphism (SNP) calling and the population genetic measures nucleotide diversity π and population differentiation F_ST_, **(e)** estimation of strain numbers by gene version reconstruction, and **(f)** differences in gene content among symbiont strains using read coverage information.

Heterogeneity in symbiont populations of individual mussels was 1 to 3 SNPs/kbp in the core genome, defined as the set of genes shared among the symbionts from all vent fields, and 5 to 11 SNPs/kbp in entire genome bins (Fig. S1, Extended Data Fig. 1). Heterogeneity was remarkably consistent in symbiont populations of different mussel individuals from the same vent field, but differed considerably between fields.

This variability is surprising, as genome-wide polymorphism rates of other sulfur-oxidizing intracellular symbionts from *Solemya* clams and *Ridgeia* tubeworms, which were also sequenced with Illumina, were an order of magnitude lower than in *Bathymodiolus* (Extended Data Tab. 2). The *Bathymodiolus* SOX symbionts had polymorphism rates more similar to those of human gut bacteria, which are 7-18 SNPs/kbp in individual microbial species within single host individuals^13^. This similarity is unexpected as in contrast to the SOX symbiont, most human gut microbes are extracellular, have a heterotrophic metabolism and frequently come into contact with a myriad of diverse microorganisms and bacteriophages within the gut, promoting rampant gene exchange^31,32^. The polymorphism rates in the SOX symbionts were also of the same order of magnitude as those observed in subpopulations of *Prochlorococcus*, the most abundant free-living bacterium in the ocean^12,33^.

### Population genomic insights into transmission and infection

The manner in which symbionts are transmitted can affect their heterogeneity, with vertically transmitted symbionts often displaying less heterogeneity than symbionts that are acquired horizontally^34^. Consistent with our findings of extensive SNP heterogeneity, symbiont nucleotide diversity π was 10 to 100 times higher in single *Bathymodiolus* mussels compared to *Solemya* clams^35^. Unlike *Solemya* symbionts that are predominantly vertically transmitted, there is reasonable evidence that *Bathymodiolus* juveniles acquire their symbionts horizontally^36,25,37,27,28^. However, it is unclear whether *Bathymodiolus* symbionts are taken up only during a permissive window early in the mussels' development, or throughout their lifetime^38^. Horizontally transmitted symbionts acquired only during a short developmental period, similar to *Ridgeia* tubeworms, would be subjected to a stronger bottleneck event than if they were continuously acquired^39^. Assuming genetic heterogeneity in the free-living stage of symbionts, within individual hosts symbiont populations would be isolated from each other, reminiscent of population dynamics in vertically transmitted symbionts (Extended Data Fig. 2). To test if this is the case, we compared the nucleotide diversity of the core genome within host individuals (π_within_) to that between hosts (pairwise, π_between_). Principal component analysis (PCA) and a PERMANOVA test on pairwise Bray-Curtis dissimilarities comparing π_within_ to π_between_ revealed that there was no significant difference between π values of hosts from the same vent field, whereas π_within_ differed significantly between vent fields (Fig. 2, Tab. S2, S3, S4, Extended Data Fig. 3). This suggests fully intermixed symbiont populations among co-occurring hosts (Fig. 2, Extended Data Fig. 2). Moreover, the fixation index (F_ST_), a measure of population differentiation^40,41^ expressed as values between 0 (no differentiation) and 1 (complete differentiation), was mostly low within a vent field (0.04-0.24) (Fig. 2, Extended Data Fig. 4). This genetic homogeneity across symbiont populations from the same vent field supports a model of intermixed symbiont populations. Alltogether, our nucleotide diversity analyses thus indicate that *Bathymodiolus* symbionts are continuously acquired from the environment throughout the host’s lifetime, confirming an earlier study based on morphological observations of continuous symbiont uptake in *Bathymodiolus*^38^.

**Figure 2.**
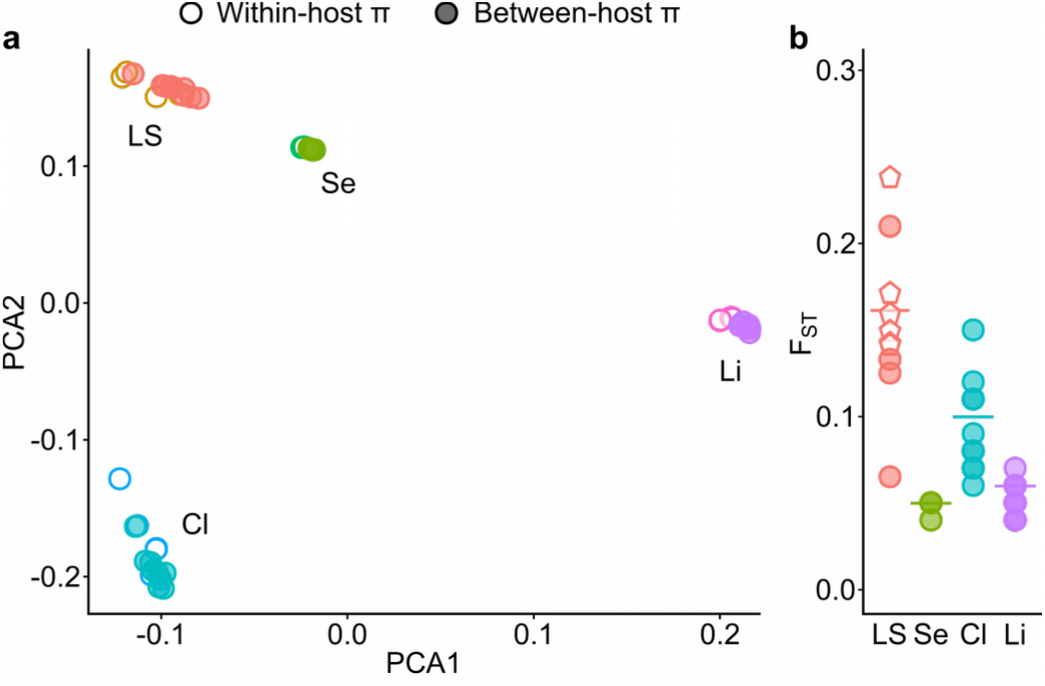
Population genetic measures π and F_ST_ show that mussels from the same site host similar symbiont populations. **(a)** Principle component analysis (PCA) of π-values (nucleotide diversity) within and in pairwise comparison between individuals for core genes of the SOX symbiont in *B.* spp from the vent fields Lucky Strike (LS), Semenov (Se), Clueless (Cl) and Lilliput (Li). Filled circles represent pairwise π-values between two hosts; empty circles represent within-host π-values. π-values cluster according to vent field but no sub-clusters appear to separate within- and between-host π-values. This is confirmed by a PERMANOVA analysis on pairwise Bray-Curtis dissimilarities: no significant difference (Pseudo-F < 1.5, P > 0.2) between within-host and pairwise between-host π; significant difference between within-host π among fields (Pseudo-F > 85, Pr < 0.001) (see Tab. S3, S4). **(b)** F_ST_-values: pairwise (symbols) and mean (line) across all host individuals per site. For vent site LS, circles represent host pairs from the same vent field, pentagonal symbols represent host pairs from two different sampling sites that are separated by approx. 150 m. At the LS vent field, a few mussel pairs showed elevated F_ST_-values, which could be explained by environmental differences between the two collection sites (discussed in Supplement).

### Symbiont strains co-exist in single host individuals

Understanding the true level of strain diversity in natural populations is a fundamental challenge in microbial ecology. To quantify strains, SNPs must be linked across genes or, if possible, entire genomes. The most sensitive ‘marker gene’ for resolving strain variability is the one that evolves most rapidly, but this is unlikely to be the same gene in all natural populations^42^. Therefore, we consider each distinct sequence of any coding gene to represent a different strain. We used more than 200 gammaproteobacterial single-copy marker genes to determine the maximum number of versions of each of these 200 genes, in each metagenome. Furthermore, we also analyzed all genes that had coverages similar to those of these single-copy marker genes, and were therefore likely present in all strains within the population. We considered a single well-supported SNP sufficient to distinguish different strains (see Methods and Supplement section 1.5).

Both approaches produced similar results, detecting up to 16 versions of the most variable symbiont genes within single mussel individuals (Fig. 3, Extended Data Fig. 5). To investigate whether sequencing depth influenced estimated strain numbers, we repeated our analyses after down-sampling the reads to the lowest coverage found in our libraries (100x; Tab. S1). This reduced the estimated strain numbers to 4-9 per host individual, showing that read coverage influenced our results (Fig. 3).

**Figure 3.**
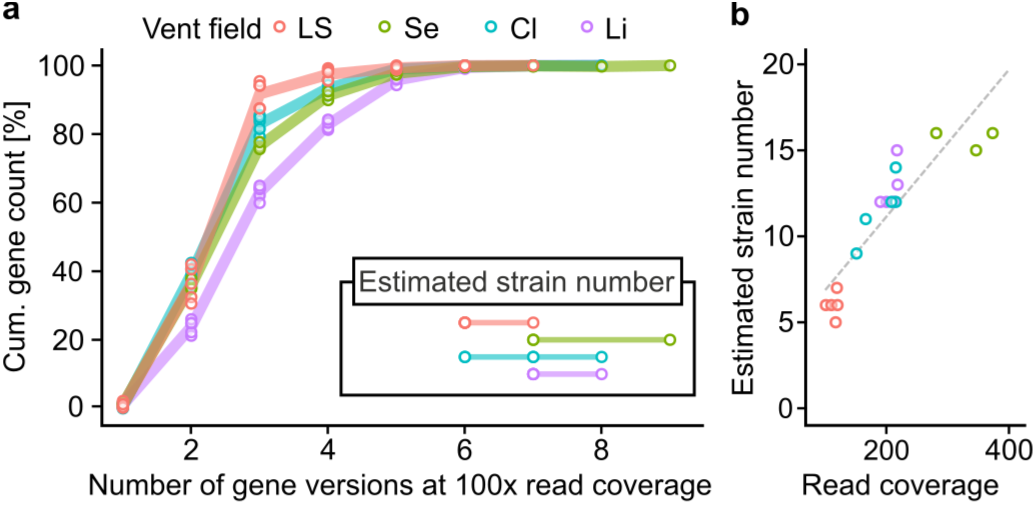
Gene version reconstruction reveals up to 16 co-occurring SOX symbiont strains in individual *Bathymodiolus* mussels. **(a)** Cumulative count shows how many genes resulted in a specific number of reconstructed gene versions. This was performed for a set of 584 to 941 genes that had a read coverage within the coverage range of gammaproteobacterial marker genes, indicating that each strain in the population encoded these. Each line represents the average cumulative gene counts across all individuals from a site and each circle represents the gene count of a single individual. These plots reveal the spectrum of variability in SOX symbiont genomes - for each gene, there were between 1 and 9 different versions in populations of single host individuals at a read coverage of 100x. The gene with the most variation, and therefore the most versions, gives the most sensitive estimate for the number of strains that may co-exist in one mussel individual. Estimates of the number of co-existing strains are shown in the inset - these are ranges of estimates, derived from the maximum number of gene versions for each host individual. **(b)** Strain numbers were estimated with full read coverage ranging from 100 to 370x, revealing that up to 16 strains can co-exist in a single host individual. The sensitivity of strain detection correlates with read coverage (spearman correlation: rs = 96, p = 4×10^−10^). LS: Lucky Strike (*B*. *azoricus),* Se: Semenov (*B. puteoserpentis),* Cl: Clueless (*B*. sp.), Li: Lilliput (*B*. sp.).

We validated our approach for estimating strain numbers by analyzing a test dataset with simulated reads from 10 published *Escherichia coli* strains with 1% genetic heterogeneity, similar to that of the *Bathymodiolus* symbionts (Tab. S5). In this test dataset, read coverage also affected estimated strain numbers: these were underestimated at 100x coverage but were closest to accurate numbers at 300x coverage (see Supplement section 2.2; Fig. S2). Our estimate of 16 co-occurring SOX strains, from a library with 373x coverage, is therefore likely realistic. We could further confirm the accuracy of our approach with long PacBio reads of a *B.* sp. individual sampled at the vent field Wideawake. We detected a maximum number of 11 distinct contigs containing the same single-copy gene, which was similar to the 12 strains we estimated using Illumina reads from the same individual (Fig. 4). Taken together, these analyses support our conclusion that, at the very least 4 to 9, but as many as 16 symbiont strains co-occured within single *Bathymodiolus* individuals. These results are surprising, as a very low level of symbiont diversity was previously assumed to be typical for these hosts based on commonly used marker genes^22,23^.

**Figure 4.**
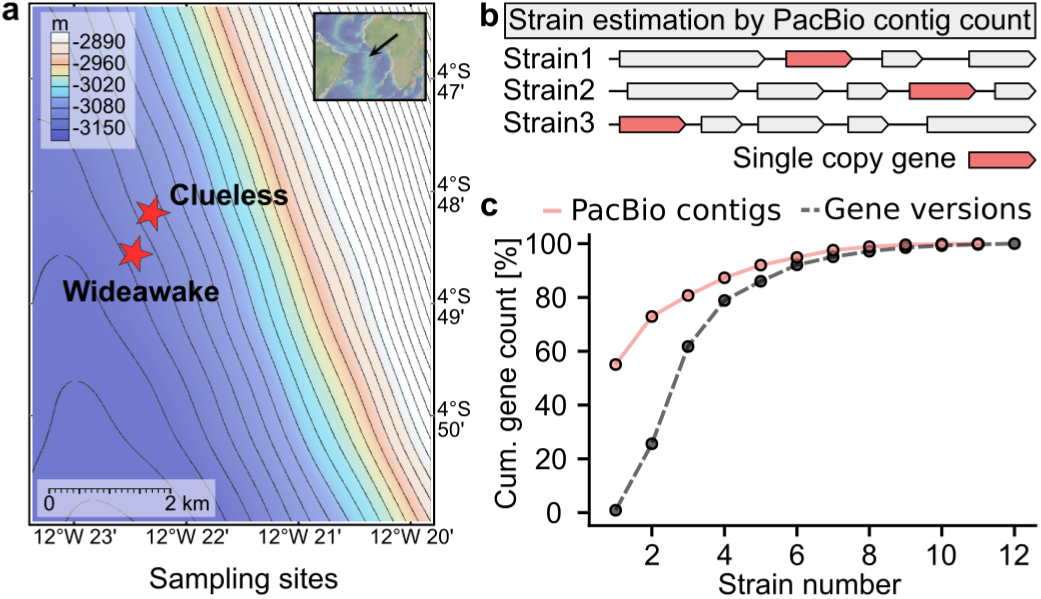
Strain number estimates from PacBio sequencing confirms strain estimation approach from gene version reconstruction of Illumina sequences. **(a)** To verify our strain number estimation workflow, we obtained long read PacBio sequences and Illumina sequences from a single B. sp. individual from the Wideawake vent field (730 m from Clueless). **(b)** PacBio sequences revealed genome rearrangements around phylogenetic marker genes. **(c)** PacBio contigs and Illumina gene version reconstructions result in similar estimates of 11 and 12 strains, respectively. Continuous red line: number of PacBio contigs containing the same single-copy genes, dashed line: number of gene versions based on Illumina sequences and plotted as in Fig. 3.

### From the pangenome to the environment: Habitat chemistry drives symbiont genome heterogeneity

Understanding the geochemical environment experienced by deep-sea organisms is challenging. In addition, the relative availability of potential energy sources can be more important than absolute availability in determining which microbial energy-generating processes are most favorable^43^. We compared symbionts from vent fields with different environmental conditions, an ideal natural experiment for investigating potential links between strain diversity and the environment. We developed a bioinformatic pipeline that used metagenomic read coverage to identify differences in gene content among co-occurring strains in our dataset of four host species from geographically and geochemically distinct vent fields. Due to uneven DNA replication rates across the entire genome, even single-copy genes encoded by all strains have a range of coverages in metagenomes^44^. To define this range, we calculated the coverage of known, single-copy gammaproteobacterial genes in each metagenome (Fig. S3). Genes with coverage values below this range were likely only encoded by a subset of the population, and were thus considered strain-specific.

Between 30 and 50% of all genes in symbiont populations from individual mussels were potentially strain-specific, indicating massive differences in the gene contents of co-occurring strains (Extended Data Tab. 3). The functions of proteins encoded by the strain-specific genes differed markedly between the four vent fields, but within a field, these were mostly consistent among host individuals (Tab. S6, Fig. 5). With few exceptions, all strain-specific genes with annotated functions could also be detected in metatranscriptomes, suggesting that differences in gene content between different strains resulted in functional differences that likely influence the fitness of symbionts and host (see Supplement section 2.3 for details, Tab. S6).

**Figure 5.**
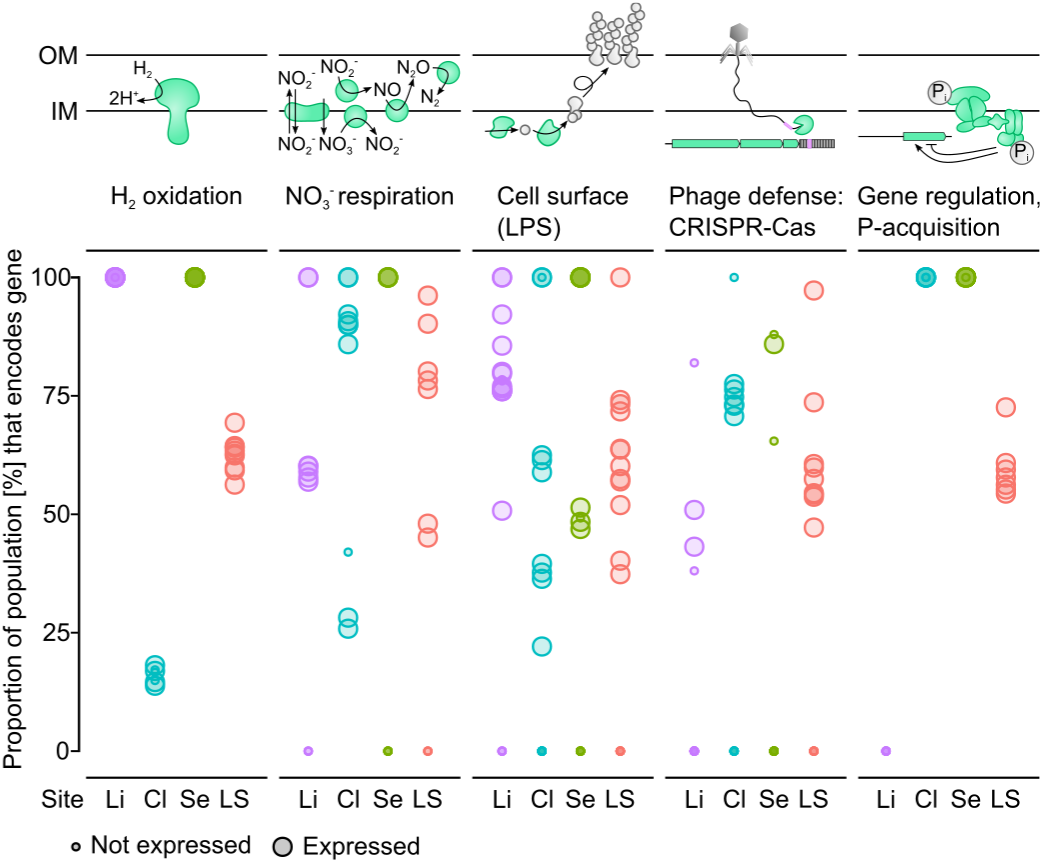
Strain-specific genes encode key functions in SOX symbionts, including energy production and interactions with hosts and phages. The proportion of strains encoding these functions was different at each hydrothermal vent field. Large dots represent single genes that were detected in the transcriptomes and small dots represent genes that were not detectable in the transcriptomes. If the capability to perform a particular function was encoded by multiple genes, then it will have multiple points, e.g. hydrogen oxidation was encoded in a cluster containing multiple genes (see Tab. S6 for more details). The proportion of a population encoding each function was calculated as the average mean coverage (3 host individuals from Semenov and 5 host individuals for each of the other vent fields) compared to the mean coverage of genes encoded by the entire population (see Materials and Methods for details). If a gene had a coverage of 0, the gene was not encoded in the symbiont genome. Colors correspond to the four different sampling sites. Li: Lilliput, Cl: Clueless, Se: Semenov, LS: Lucky Strike.

More than 80% of the strain-specific genes encoded hypothetical proteins with unknown functions. Remarkably, although only a small proportion of the strain-specific genes could be annotated, these genes encode proteins involved in key functions such as synthesis of cell-surface components, environmental phosphate (P_i_) sensing and acquisition, cell-cell interactions and phage defense (Fig. 5, Extended Data Fig. 6, Extended Data Fig. 7, Tab. S6). Hydrogen oxidation and nitrate reduction genes were also strain-specific in Mid-Atlantic Ridge populations, as shown previously in *B. septemdierum* from the West Pacific (Fig. 5)^29^. Some of the strain-specific symbiont genes may provide a selective advantage depending on the vent environment. For example, all mussels from vent fields with the highest hydrogen concentrations had a larger proportion of strains encoding hydrogenases, and those from fields with the lowest concentrations had the smallest proportion of strains encoding these enzymes (see Supplement 2.3). Ikuta *et al*.^29^ also found differences in the relative proportions of strains that could oxidize hydrogen in a single *Bathymodiolus* species sampled from two vents. However, as most individuals sampled from one field were small juveniles, and most collected from the second field were adults, it was unclear whether this reflected site-specific differences in hydrogen availability, or changes during host development.

Genes involved in phosphate metabolism were another example of strain-specific variability that could provide a selective advantage depending on vent conditions. These genes were in a single cluster and encoded the high-affinity phosphate transport system PstSCAB, the regulatory protein PhoU and the two-component regulatory system PhoR-PhoB^45^. In addition to phosphorous metabolism, PhoR-B can also affect other functions such as secondary metabolite production and virulence^46–48^. Considering the key role of these genes in cellular metabolism, it is surprising that this gene cluster was only encoded by the entire population of symbiont strains in mussels from two vent fields (Fig. 5). These genes were not present in any of the symbiont strains from Lilliput mussels, and only in some symbiont strains of Lucky Strike mussels, as confirmed by read mapping against symbiont bins (Fig. 5). At most hydrothermal vents P_i_ concentrations are unknown. However, soluble P_i_ depends on iron concentrations, which are reported to vary substantially between vent fields, raising the possibility that environmental P_i_ availability drives the loss or gain of P_i_-related genes in symbiont populations (see Supplement section 2.3). Genes involved in P_i_ uptake and regulation were also strain-specific in *Prochlorococcus,* and their presence was linked to environmental P_i_ concentrations^49,50^. The SOX symbionts of vesicoymid clams and free-living relatives *Thioglobus* spp. from the SUP05 clade appear to also lack the PstSCAB genes based on our analyses of their published genomes (accession numbers of symbionts: JARW01000002, DDCF01000009, NC_009465, NC_008610; SUP05: CP010552, CP006911, CP008725, GG729964). However, to our knowledge no other bacteria have been described to miss both, the PhoR-B and PstSCAB systems. The symbiotic and free-living SOX bacteria that lack PstSCAB might use a low-affinity P_i_-transporter to acquire P_i_, as genes for these transporters were encoded in all of the analyzed genomes.

Oxygen concentrations fluctuate at vents due to dynamic mixing of anoxic hydrothermal fluids and oxygen-rich deep-sea seawater, and accordingly, mussel symbionts can use alternative electron acceptors such as nitrate^51–54^. Complete reduction of nitrate to dinitrogen gas (N_2_) requires four enzymes: respiratory nitrate reductase (Nar), nitrite reductase (Nir), nitric oxide reductase (Nor) and nitrous oxide reductase (Nos)^55^. In contrast to the genes needed for oxygen respiration, which were present in all symbiont populations, the prevalence of genes encoding all four steps of nitrate reduction to N_2_ was highly variable among vent fields, among mussels from the same field, and even within symbiont populations of single mussels (Fig. 5, Extended Data Fig. 6). For example, in Lucky Strike individuals, the enzyme for the reduction of nitrate to nitrous oxide (N_2_O) was encoded by 30 to 100% of the population, whereas the ability to perform the last step from N_2_O to N_2_ was not encoded at all. These variable abundances within symbiont populations suggest that each of the three steps of nitrate reduction to N_2_O might be performed by a different subset of strains within a single host (Fig. 5, Extended Data Fig. 6). The remarkable modularity of nitrate respiration genes in *Bathymodiolus* symbionts, as well as in other symbiotic and free-living bacteria^56,57^, suggests that these genes are particularly prone to loss and gain. This raises the intriguing possibility that intricate interactions between microbes exchanging N intermediates are widespread in natural populations. Such a ‘division of labor’ may be beneficial as it could increase community nutrient consumption and avoid accumulation of toxic intermediates^58,59^. In fact, when individual reactions of the denitrification pathway are subdivided among strains, the toxicity of the intermediates might result in dependence between strains producing and strains consuming toxic intermediates.

Our results revealed that *Bathymodiolus* SOX symbionts have pangenomes with considerable functional diversity among co-existing strains, and this metabolic diversity may be linked to vent geochemistry. Together with our findings of continuous uptake of symbionts throughout the mussel's lifetime, these results suggest constant symbiont strain shuffling between the environment and host as well as among co-occurring hosts. Given that the *Bathymodiolus* SOX symbiosis has evolved multiple times in convergent evolution from within the SUP05 clade^24^, it is possible that some of this strain reshuffling might also involve free-living SUP05 bacteria. This would allow *Bathymodiolus* to associate with those strains that are best adapted to the vent environment, even within the lifetime of an individual mussel. Rapid reshuffling of microbes has also been observed in other systems such as the human gut microbiota, where food intake has a direct and immediate effect on the microbial community^60^. At hydrothermal vents, exchange of symbiont strains would result in rapid holobiont adaptation to local conditions within the lifetime of individual mussel hosts. Such genomic flexibility of the symbionts may underpin the productivity and global success of *Bathymodiolus* mussels in these ecosystems.

### A new model of evolutionary stability for one-to-many symbioses

Ecological theory predicts that if two different organisms share a limited resource, one will out-compete the other, unless mechanisms such as niche partitioning allow their stable co-existence^61,62^. Can these theories explain our results of co-existing strains, and how strain diversity and competition impact symbiosis stability? If competing symbionts differ in the net mutualistic benefit they provide, hosts can benefit by evolving mechanisms to differentially distribute costly resources to their partners. This can drive the evolution of specialized structures, such as compartments, with low symbiont diversity. In these cases, discrimination by hosts is important because a costly resource, such as photosynthate in the legume-*Rhizobia* symbioses, is provided^63^. But what if the symbiosis has low costs to the host?

Knowledge of costs and benefits of symbiotic associations is central to understanding their evolutionary trajectories. Beyond nutritional benefits gained through symbiont digestion^64^, the benefits as well as the costs for *Bathymodiolus* mussels have not been extensively investigated. Possible costs include maintaining host-symbiont recognition mechanisms, transporting symbiont substrates into bacteriocyte vacuoles and waste products out, and dealing with toxic reactive oxygen species produced by symbiont metabolism. We currently understand even less about the costs and benefits for the symbionts. They may benefit from improved access to reduced and oxidized substrates, such as sulfide and oxygen, which often do not overlap spatially or temporally (although see^65^). Costs possibly include maintenance of recognition and intracellular survival mechanisms, and loss of a substantial part of the population through intracellular digestion.

In contrast to many well-characterized symbioses (e.g. see^63^), there is one substantial cost that *Bathymodiolus* does not have to bear - the cost of ‘feeding’ its symbionts. This is because the symbionts’ major energy sources come from the vent environment. *Bathymodiolus* symbioses therefore more closely resemble byproduct mutualisms, which are considered ‘low-to-no-cost’ associations^66^. Such lower costs for the host would shift the cost-benefit balance so that a greater range of symbiont strains with distinct metabolic capabilities could still provide a net benefit to the host. This implies that ‘low-quality’ symbionts that grow more slowly could thus co-occur alongside high-quality symbionts. Moreover, strain diversity has additional ecological and evolutionary benefits such as bacterial protection against bacteriophage attacks, and holobiont adaptation to new and changing environments. A ‘low-quality’ symbiont under certain conditions can become a ‘high-quality’ symbiont when environmental conditions change^67^.

Low costs can also remove potential incentives for partners to ‘cheat’^68^. ‘Cheating’ is defined as using services provided by the host, and providing fewer or no services in return^66^. In the case of *Bathymodiolus*, the host would appear fully in control of the transfer of benefits from symbionts to the host. Regardless of whether symbionts share the products of carbon fixation immediately with their hosts through ‘leaking’ of small compounds, or whether these are primarily directed towards symbiont cell biosynthesis, intracellular digestion of symbiont cells ensures that all the products of symbiont primary production are eventually transferred to the host. Furthermore, because the symbionts gain the bulk of their energy from the environment, instead of destabilizing the association as described by current evolutionary models, competition between different symbiont types could be beneficial for the host, if it results in the dominance of strains that more effectively transform geochemical energy in the vent environment into biomass and thus into host nutrition^14^.

## Conclusion

Our view of microbial diversity has long been shaped by our limited ability to accurately assess the enormous diversity of natural communities^69^. Metagenomics is rapidly changing this view, revealing that strain diversity has been vastly underestimated. Our study shows that strain diversity is pervasive in the sulfur-oxidizing symbionts of *Bathymodiolus* mussels, and that this diversity, invisible at the level of marker genes, has massive genome-wide effects. Symbioses between corals and their intracellular photosynthetic algae are another prominent example where strain diversity may be common, although it is still unclear how much of this diversity is due to different gene copies within a single eukaryotic genome^15,70,71^. High symbiont diversity was also recently identified in the photosynthetic symbionts of marine protists^72^.

This unexpected diversity has wide-ranging implications for the function and evolution of host-microbe associations. Despite this, it is currently not considered by most evolutionary theories, because these theories have been shaped by decades of study focused on models of symbiosis in which the host bears the enormous cost of ‘feeding’ the symbionts, and symbiont genetic diversity is highly restricted. We provide a new theoretical framework that could explain the unexpected prevalence and evolutionary stability of strain diversity in beneficial host-microbe associations, where the environment provides for the symbionts’ nutrition. This is the case for a diverse range of host-microbe associations from marine chemosynthetic and photosynthetic symbioses to the human digestive tract. Considering the substantial evidence that biodiversity underpins ecosystem stability, productivity, and resistance to invasion and parasitism^4^, we predict that strain variation should be widespread in ‘low-cost’ associations such as these. Clearly, new concepts are needed that extend evolutionary theories that were developed based on earlier studies of beneficial associations to a more united framework that can explain the wide range of host – microbe associations recent research is unveiling.

## Methods

### Sample collection

Four *Bathymodiolus* species from four vent fields were collected during three research cruises at hydrothermal vents along the Mid-Atlantic Ridge (MAR). Mussels from the same vent field belonged to the same host species based on their mitochondrial cytochrome c oxidase subunit I sequences. Symbiont-containing gill tissues were dissected from five mussel individuals from each of the following vent fields: Lucky Strike (site ‘Montsegur' 37°17'19.1760"N, 32°16'32.0520"W; site ‘Eiffel Tower’ 37°17'20.8320"N, 32°16'31.7640"W), Clueless (4°48'11.7594"S, 12°22'18.4814"W) and Lilliput (9°32'47.6412"S, 13°12'35.0388"W). From these fields, samples were always dissected from the middle of each gill. From the Semenov-2 field, gill pieces were dissected from the gill edges of three individuals (location ‘Ash Lighthouse’ 13°30'48.4812"N, 44°57'47.2788"W). One additional mussel individual was sampled at Wideawake (4°48'37.5599"S, 12°22'20.5201"W, 730 m from Clueless). From this individual, the whole gill was homogenized in a Dounce tissue grinder (Sigma, Germany) and a subsample used for DNA sequencing. For an overview of these locations and samples, see the map in Fig. 1 and Tab. S7. Gill tissue pieces were either frozen directly at −80 °C or fixed in RNAlater according to the manufacturer’s instructions (Sigma, Germany) and subsequently frozen at −80 °C.

### Nucleic acid extraction and metagenome sequencing

DNA was extracted from gill pieces with commercially available kits (Tab. S8). RNA was extracted using the AllPrep kit (Tab. S9, Qiagen, Germany). From the symbiont homogenate from Wideawake, DNA was extracted according to Zhou *et al*.^73^. For each vent field, one reference SOX symbiont bin was produced from co-assemblies of metagenomes from multiple individuals as follows (see Tab. S1 for reference genome statistics). Metagenomes were sequenced with Illumina or PacBio technology (see Supplement section 1.1 for details). Metagenomes were assembled from Illumina reads using IDBA-ud (v 1.1.1)^74^ and SPAdes (v 3.2.2)^75^, and genome bins were produced using a custom combination of differential coverage analysis with GBtools (v 2.4.5)^76^ and contig connectivity analysis^77^, and annotated with RASTtk^78^ (see Supplement section 1.2 for details).

### Transcriptome sequencing and analysis

Transcriptome reads were mapped to reference genomes with BBMap (v 36.x, Bushnell B. - BBMap - sourceforge.net/projects/bbmap/). The number of transcripts per gene was estimated with featureCounts^79^. Transcripts were normalized for different sequencing depths across libraries and for the gene length using edgeR with trimmed mean of M values (TMM) normalization^80,81^.

### SNP calling and population structure analysis

SNPs were called from reads of each individual sample mapped to the consensus symbiont bin and filtered, both performed with the Genome Analysis Toolkit (*GATK* v3.3.0; see Supplement section 1.3 for details)^82^. Rather than using the default settings for diploid genomes, we chose a ploidy setting of 10, as this better reflects a mixture of coexisting bacterial strains (Tab. S10). The symbiont population structure within and between host individuals was investigated by calculating nucleotide diversity π and the fixation index F_ST_ based on SNP frequencies (see Supplement section 1.6 for details)^13^.

### Core genome calculation and detection of strain-specific genes

We developed a bioinformatic pipeline to identify strain-specific genes in metagenomes based on relative read coverage (see Supplement section 1.4 for details). Briefly, we defined the coverage range of genes that are encoded by each strain in the population, based on single-copy gammaproteobacterial marker genes^83^, and regarded all genes with coverage below this range as potentially strain specific. For some of these, multiple gene copies (coding sequences with the same annotation) were present in one metagenome. We excluded these genes from further analyses because it is possible that all strains encoded these, but that rearrangements led to different gene neighborhoods, causing these genes to fall on different contigs in the genome assemblies.

### Strain number estimation and test simulation

We estimated the number of strains by using the number of gene sequence versions that could be reconstructed based on SNP linkage and frequency as a proxy. These distinct sequence versions were reconstructed for gammaproteobacterial marker genes from PhylaAmphora^83^, as well as for all the genes encoded by each strain in the symbiont population in a single mussel (identified by read coverage, see above) using the tool ViQuaS (v 1.3)^84^. We created a test dataset with parameters that were similar to the sequencing data used in this study, by simulating Illumina reads from 10 publicly available *E. coli* genomes with ART (v 2.5.8)^85^ (Tab. S5). Reads were pooled in even and uneven ratios to simulate different abundance patterns of strains in the population. Both datasets were analyzed with our strain estimation pipeline for two coverage depths 100x and 300x (see Supplement section 1.5 for details).

### Code and data availability

Custom code is available on the github repository https://github.com/rbcan/MARsym_paper for detailed information of the computing steps. All sequencing reads and symbiont bins used in this study can be found at ENA under the accession number PRJEB28154.

## Acknowledgements

We thank the captains, crews and ROV teams on the cruises BioBaz (2013), ODEMAR (2014), M78-2 (2009), Atalante Cruise Leg – 2 (2008) on board of the research vessels Pourquoi Pas?, FS Meteor and L'Atalante and the chief scientists François Lallier, Javie Excartin and Muriel Andreani, Richard Seifert and Colin Devey. Thank you to Adrien Assié, Christian Borowski, Corinna Breusing and Karina van der Heijden for sample treatment and fixation on board, and Målin Tietjen for the extraction of RNA from the samples of vent fields Semenov, Clueless and Lilliput. We also thank Christian Quast and Hanno Teeling for technical support. We thank Tal Dagan for the discussions and input during the project and on the written manuscript.

This study was funded by the Max Planck Society, the MARUM DFG-Research Center / Excellence Cluster “The Ocean in the Earth System” at the University of Bremen, the DFG CRC 1182 "Origin and Function of Metaorganisms", the German Research Foundation (RV Meteor M78-2 cruise), an ERC Advanced Grant (BathyBiome, 340535), and a Gordon and Betty Moore Foundation Marine Microbial Initiative Investigator Award to ND (Grant GBMF3811).

## Contributions

R.A., J.P., L.S. and N.D conceived the study. R.A. and J.P. wrote the manuscript, with support from N.D., and contributions and revisions from all other co-authors. R.A. developed the metagenomic workflow for polymorphism detection, strain reconstruction, identification of strain-specific genes and analyzed the data with the exceptions described in the following. S.R. conducted the core-genome calculation, read simulation analyses, provided support for the statistical analyses and drafted respective manuscript sections. L.S. extracted nucleic acids for samples from Lucky Strike, Semenov and Wideawake, and conducted and evaluated the PacBio assembly. A.K. developed and provided an R-script for the calculation of π and F_ST_. H.T. sequenced metagenomes from vent fields Clueless and Lilliput.

## References

1. Pankey, M. S. et al. Host-selected mutations converging on a global regulator drive an adaptive leap towards symbiosis in bacteria. eLife 6, e24414 (2017).

2. Viana, D. et al. A single natural nucleotide mutation alters bacterial pathogen host-tropism. Nat. Genet. 47, 361–366 (2015).

3. Greenblum, S., Carr, R. & Borenstein, E. Extensive strain-level copy-number variation across human gut microbiome species. Cell 160, 583–594 (2015).

4. Tilman, D., Isbell, F. & Cowles, J. M. Biodiversity and ecosystem functioning. Annu. Rev. Ecol. Evol. Syst. 45, 471–493 (2014).

5. Hooper D. U. et al. Effects of biodiversity on ecosystem functioning: a consensus of current knowledge. Ecol. Monogr. 75, 3–35 (2005).

6. Frank, S. A. Host–symbiont conflict over the mixing of symbiotic lineages. Proc R Soc Lond B 263, 339–344 (1996).

7. Sachs, J. L. et al. Host control over infection and proliferation of a cheater symbiont. J. Evol. Biol. 23, 1919–1927 (2010).

8. Bulgheresi, S. et al. A new C-type lectin similar to the human immunoreceptor DC-SIGN mediates symbiont acquisition by a marine nematode. Appl. Environ. Microbiol. 72, 2950–2956 (2006).

9. Nyholm, S. V. & McFall-Ngai, M. The winnowing: establishing the squid–vibrio symbiosis. Nat. Rev. Microbiol. 2, 632–642 (2004).

10. Delmont, T. O. & Eren, A. M. Linking pangenomes and metagenomes: the Prochlorococcus metapangenome. PeerJ 6, e4320 (2018).

11. Engel, P., Stepanauskas, R. & Moran, N. A. Hidden diversity in honey bee gut symbionts detected by single-cell genomics. PLoS Genet. 10, e1004596 (2014).

12. Kashtan, N. et al. Single-cell genomics reveals hundreds of coexisting subpopulations in wild Prochlorococcus. Science 344, 416–420 (2014).

13. Schloissnig, S. et al. Genomic variation landscape of the human gut microbiome. Nature 493, 45–50 (2013).

14. Batstone, R. T., Carscadden, K. A., Afkhami, M. E. & Frederickson, M. E. Using niche breadth theory to explain generalization in mutualisms. Ecology 99, 1039–1050 (2018).

15. Rowan, R. & Knowlton, N. Intraspecific diversity and ecological zonation in coral-algal symbiosis. Proc. Natl. Acad. Sci. 92, 2850–2853 (1995).

16. Foster, K. R., Schluter, J., Coyte, K. Z. & Rakoff-Nahoum, S. The evolution of the host microbiome as an ecosystem on a leash. Nature 548, 43–51 (2017).

17. Bongrand, C. et al. A genomic comparison of 13 symbiotic Vibrio fischeri isolates from the perspective of their host source and colonization behavior. ISME J. 10, 2907–2917 (2016).

18. Quince, C. et al. DESMAN: a new tool for de novo extraction of strains from metagenomes. Genome Biol. 18, 181 (2017).

19. Cleary, B. et al. Detection of low-abundance bacterial strains in metagenomic datasets by eigengenome partitioning. Nat. Biotechnol. 33, 1053–1060 (2015).

20. Nielsen, H. B. et al. Identification and assembly of genomes and genetic elements in complex metagenomic samples without using reference genomes. Nat. Biotechnol. 32, 822–828 (2014).

21. Petersen, J. M. & Dubilier, N. Methanotrophic symbioses in marine invertebrates. Environ. Microbiol. Rep. 1, 319–335 (2009).

22. Dubilier, N., Bergin, C. & Lott, C. Symbiotic diversity in marine animals: the art of harnessing chemosynthesis. Nat. Rev. Microbiol. 6, 725–740 (2008).

23. Duperron, S. et al. A dual symbiosis shared by two mussel species, Bathymodiolus azoricus and Bathymodiolus puteoserpentis (Bivalvia: Mytilidae), from hydrothermal vents along the northern Mid-Atlantic Ridge. Environ. Microbiol. 8, 1441–1447 (2006).

24. Petersen, J. M., Wentrup, C., Verna, C., Knittel, K. & Dubilier, N. Origins and evolutionary flexibility of chemosynthetic symbionts from deep-sea animals. Biol. Bull. 223, 123–137 (2012).

25. Duperron, S. The diversity of deep-sea mussels and their bacterial symbioses. in The Vent and Seep Biota (ed. Kiel, S.) 33, 137–167 (Springer Netherlands, 2010).

26. Duperron, S. et al. Diversity, relative abundance and metabolic potential of bacterial endosymbionts in three Bathymodiolus mussel species from cold seeps in the Gulf of Mexico. Environ. Microbiol. 9, 1423–1438 (2007).

27. DeChaine, E. G. & Cavanaugh, C. M. Symbioses of methanotrophs and deep-sea mussels (Mytilidae: Bathymodiolinae). in Molecular Basis of Symbiosis (ed. Overmann, P. D. J.) 227–249 (Springer Berlin Heidelberg, 2005).

28. Won, Y.-J. et al. Environmental acquisition of thiotrophic endosymbionts by deep-sea mussels of the genus Bathymodiolus. Appl. Environ. Microbiol. 69, 6785–6792 (2003).

29. Ikuta, T. et al. Heterogeneous composition of key metabolic gene clusters in a vent mussel symbiont population. ISME J. 10, 990–1001 (2016).

30. Heath, K. D. & Stinchcombe, J. R. Explaining mutualism variation: a new evolutionary paradox? Evolution 68, 309–317 (2014).

31. Smillie, C. S. et al. Ecology drives a global network of gene exchange connecting the human microbiome. Nature 480, 241–244 (2011).

32. Hehemann, J.-H. et al. Transfer of carbohydrate-active enzymes from marine bacteria to Japanese gut microbiota. Nature 464, 908–912 (2010).

33. McInerney, J. O., McNally, A. & O’Connell, M. J. Why prokaryotes have pangenomes. Nat. Microbiol. 2, 17040 (2017).

34. Douglas, A. E. The ecology of symbiotic micro-organisms. in Advances in Ecological Research (eds. Begon, M. & Fitter, A. H.) 26, 69–103 (Academic Press, 1995).

35. Russell, S. L., Corbett-Detig, R. B. & Cavanaugh, C. M. Mixed transmission modes and dynamic genome evolution in an obligate animal–bacterial symbiosis. ISME J. 11, 1359 (2017).

36. Laming, S. R., Duperron, S., Cunha, M. R. & Gaudron, S. M. Settled, symbiotic, then sexually mature: adaptive developmental anatomy in the deep-sea, chemosymbiotic mussel Idas modiolaeformis. Mar. Biol. 161, 1319–1333 (2014).

37. Won, Y.-J., Jones, W. J. & Vrijenhoek, R. C. Absence of cospeciation between deep-sea Mytilids and their thiotrophic endosymbionts. J. Shellfish Res. 27, 129–138 (2008).

38. Wentrup, C., Wendeberg, A., Schimak, M., Borowski, C. & Dubilier, N. Forever competent: deep-sea bivalves are colonized by their chemosynthetic symbionts throughout their lifetime. Environ. Microbiol. 16, 3699–3713 (2014).

39. Chaston, J. & Goodrich-Blair, H. Common trends in mutualism revealed by model associations between invertebrates and bacteria. FEMS Microbiol. Rev. 34, 41–58 (2010).

40. Wright, S. Evolution and the genetics of populations. The theory of gene frequencies. Vol. 2, (University of Chicago Press, 1969).

41. Wright, S. Isolation by distance. Genetics 28, 114–138 (1943).

42. Lan, Y., Rosen, G. & Hershberg, R. Marker genes that are less conserved in their sequences are useful for predicting genome-wide similarity levels between closely related prokaryotic strains. Microbiome 4, 18 (2016).

43. Perner, M. et al. Linking geology, fluid chemistry, and microbial activity of basalt- and ultramafic-hosted deep-sea hydrothermal vent environments. Geobiology 11, 340–355 (2013).

44. Brown, C. T., Olm, M. R., Thomas, B. C. & Banfield, J. F. Measurement of bacterial replication rates in microbial communities. Nat. Biotechnol. 34, 1256 (2016).

45. Hsieh, Y.-J. & Wanner, B. L. Global regulation by the seven-component Pi signaling system. Curr. Opin. Microbiol. 13, 198–203 (2010).

46. Romano, S., Schulz-Vogt, H. N., González, J. M. & Bondarev, V. Phosphate limitation induces drastic physiological changes, virulence-related gene expression, and secondary metabolite production in Pseudovibrio sp. strain FO-BEG1. Appl. Environ. Microbiol. 81, 3518–3528 (2015).

47. Santos-Beneit, F. The Pho regulon: a huge regulatory network in bacteria. Front. Microbiol. 6, (2015).

48. Lamarche, M. G., Wanner, B. L., Crépin, S. & Harel, J. The phosphate regulon and bacterial virulence: a regulatory network connecting phosphate homeostasis and pathogenesis. FEMS Microbiol. Rev. 32, 461–473 (2008).

49. Martiny, A. C., Huang, Y. & Li, W. Occurrence of phosphate acquisition genes in Prochlorococcus cells from different ocean regions. Environ. Microbiol. 11, 1340–1347 (2009).

50. Martiny, A. C., Coleman, M. L. & Chisholm, S. W. Phosphate acquisition genes in Prochlorococcus ecotypes: Evidence for genome-wide adaptation. Proc. Natl. Acad. Sci. U. S. A. 103, 12552–12557 (2006).

51. Zielinski, F. U., Gennerich, H.-H., Borowski, C., Wenzhöfer, F. & Dubilier, N. In situ measurements of hydrogen sulfide, oxygen, and temperature in diffuse fluids of an ultramafic-hosted hydrothermal vent field (Logatchev, 14°45′N, Mid-Atlantic Ridge): Implications for chemosymbiotic bathymodiolin mussels. Geochem. Geophys. Geosystems 12, Q0AE04 (2011).

52. Kuwahara, H. et al. Reduced Genome of the thioautotrophic intracellular symbiont in a deep-sea clam, Calyptogena okutanii. Curr. Biol. 17, 881–886 (2007).

53. Hentschel, U., Hand, S. & Felbeck, H. The contribution of nitrate respiration to the energy budget of the symbiont-containing clam Lucinoma aequizonata: a calorimetric study. J. Exp. Biol. 199, 427–433 (1996).

54. Hentschel, U., Cary, S. C. & Felbeck, H. Nitrate respiration in chemoautotrophic symbionts of the bivalve Lucinoma aequizonata. Mar. Ecol. Prog. Ser. 94, 35–41 (1993).

55. Kraft, B., Strous, M. & Tegetmeyer, H. E. Microbial nitrate respiration – Genes, enzymes and environmental distribution. J. Biotechnol. 155, 104–117 (2011).

56. Shah, V., Chang, B. X. & Morris, R. M. Cultivation of a chemoautotroph from the SUP05 clade of marine bacteria that produces nitrite and consumes ammonium. ISME J. 11, 263–271 (2017).

57. Kleiner, M., Petersen, J. M. & Dubilier, N. Convergent and divergent evolution of metabolism in sulfur-oxidizing symbionts and the role of horizontal gene transfer. Curr. Opin. Microbiol. 15, 621–631 (2012).

58. Savage, V. M., Webb, C. T. & Norberg, J. A general multi-trait-based framework for studying the effects of biodiversity on ecosystem functioning. J. Theor. Biol. 247, 213–229 (2007).

59. Lindemann, S. R. et al. Engineering microbial consortia for controllable outputs. ISME J. 10, 2077–2084 (2016).

60. David, L. A. et al. Diet rapidly and reproducibly alters the human gut microbiome. Nature 505, 559–563 (2014).

61. Ghoul, M. & Mitri, S. The ecology and evolution of microbial competition. Trends Microbiol. 24, 833–845 (2016).

62. Hardin, G. The competitive exclusion principle. Science 131, 1292–1297 (1960).

63. Udvardi, M. & Poole, P. S. Transport and metabolism in legume-rhizobia symbioses. Annu. Rev. Plant Biol. 64, 781–805 (2013).

64. Zheng, P. et al. Insights into deep-sea adaptations and host-symbiont interactions: a comparative transcriptome study on Bathymodiolus mussels and their coastal relatives. Mol. Ecol. 26, 5133–5148 (2017).

65. Garcia, J. R. & Gerardo, N. M. The symbiont side of symbiosis: do microbes really benefit? Front. Microbiol. 5, (2014).

66. Douglas, A. E. Conflict, cheats and the persistence of symbioses. New Phytol. 177, 849–858 (2008).

67. Palmer, T. M. et al. Synergy of multiple partners, including freeloaders, increases host fitness in a multispecies mutualism. Proc. Natl. Acad. Sci. 107, 17234–17239 (2010).

68. Foster, K. R. & Wenseleers, T. A general model for the evolution of mutualisms. J. Evol. Biol. 19, 1283–1293

69. McLaren, M. R. & Callahan, B. J. In nature, there is only diversity. mBio 9, e02149–17 (2018).

70. Wooldridge Scott A. Is the coral‐algae symbiosis really ‘mutually beneficial’ for the partners? BioEssays 32, 615–625 (2010).

71. Oppen, M. J. H. van, Palstra, F. P., Piquet, A. M.-T. & Miller, D. J. Patterns of coral– dinoflagellate associations in Acropora: significance of local availability and physiology of Symbiodinium strains and host–symbiont selectivity. Proc. R. Soc. Lond. B Biol. Sci. 268, 1759–1767 (2001).

72. Brisbin, M. M., Mesrop, L. Y., Grossmann, M. M. & Mitarai, S. Intra-host symbiont diversity and extended symbiont maintenance in photosymbiotic Acantharea (clade F). Front. Microbiol. 9, (2018).

73. Zhou, J., Bruns, M. A. & Tiedje, J. M. DNA recovery from soils of diverse composition. Appl. Environ. Microbiol. 62, 316–322 (1996).

74. Peng, Y., Leung, H. C. M., Yiu, S. M. & Chin, F. Y. L. IDBA-UD: a de novo assembler for single-cell and metagenomic sequencing data with highly uneven depth. Bioinformatics 28, 1420–1428 (2012).

75. Bankevich, A. et al. SPAdes: a new genome assembly algorithm and its applications to single-cell sequencing. J. Comput. Biol. 19, 455–477 (2012).

76. Seah, B. K. B. & Gruber-Vodicka, H. R. gbtools: interactive visualization of metagenome bins in R. Front. Microbiol. 6, (2015).

77. Albertsen, M. et al. Genome sequences of rare, uncultured bacteria obtained by differential coverage binning of multiple metagenomes. Nat. Biotechnol. 31, 533–538 (2013).

78. Brettin, T. et al. RASTtk: a modular and extensible implementation of the RAST algorithm for building custom annotation pipelines and annotating batches of genomes. Sci. Rep. 5, 8365 (2015).

79. Liao, Y., Smyth, G. K. & Shi, W. featureCounts: an efficient general purpose program for assigning sequence reads to genomic features. Bioinforma. Oxf. Engl. 30, 923–930 (2014).

80. Robinson, M. D., McCarthy, D. J. & Smyth, G. K. edgeR: a Bioconductor package for differential expression analysis of digital gene expression data. Bioinformatics 26, 139–140 (2010).

81. Robinson, M. D. & Oshlack, A. A scaling normalization method for differential expression analysis of RNA-seq data. Genome Biol. 11, R25 (2010).

82. McKenna, A. et al. The Genome Analysis Toolkit: A MapReduce framework for analyzing next-generation DNA sequencing data. Genome Res. 20, 1297–1303 (2010).

83. Wang, Z. & Wu, M. A phylum-level bacterial phylogenetic marker database. Mol. Biol. Evol. 30, 1258–1262 (2013).

84. Jayasundara, D. et al. ViQuaS: an improved reconstruction pipeline for viral quasispecies spectra generated by next-generation sequencing. Bioinforma. Oxf. Engl. 31, 886–896 (2015).

85. Huang, W., Li, L., Myers, J. R. & Marth, G. T. ART: a next-generation sequencing read simulator. Bioinformatics 28, 593–594 (2012).

