## Extended Data for "Diversity matters: Deep-sea mussels harbor multiple symbiont strains"

**Extended Data Table 1 | Number of SNPs in the symbiont 16S rRNA gene within individual hosts at the four vent fields.** The position on the 16S rRNA gene, SNP frequencies and nucleotide changes are indicated in light grey boxes. The following host species correspond to listed vent fields: Lucky Strike – *B. azoricus*, Semenov – *B. puteoserpentis*, Clueless – *B.* sp., Lilliput – *B.* sp.

| | Lucky Strike | | Semenov | Clueless | | Lilliput | |
|---|---|---|---|---|---|---|---|
| Host individual 1 | 0 | | 0 | 0 | | 2* | 1021 (8%) T>A<br>1022 (8%) T>A |
| Host individual 2 | 1’ | 1018 (11%) G>A | 0 | 0 | | 2* | 1021 (8%) T>A<br>1022 (8%) T>A |
| Host individual 3 | 1’ | 1018 (16%) G>A | 0 | 0 | | 0 | |
| Host individual 4 | 2’ | 1018 (13%) G>A<br>930 (9%) C>T | - | 1 | 1036 (10%) T>C | 0 | |
| Host individual 5 | 0 | | - | 0 | | 0 | |

‘ 1 SNP is the same in all 3 individuals
* 2 SNPs are the same in both individuals

**Extended Data Table 3 | Counts and percentages of low-coverage genes in within-host symbiont populations.** Subsets represent counts that excluded all genes annotated as “hypothetical protein” and are further defined as strain-specific genes (one-copy genes with lower coverage in most or all symbiont populations from that site) and low-coverage genes with further copies in the genome. The following host species correspond to listed vent fields: Lucky Strike – *B. azoricus*, Semenov – *B. puteoserpentis*, Clueless – *B.* sp., Lilliput – *B.* sp.

| Vent site | Total low-coverage genes | w/o hypotheticals | | |
|---|---|---|---|---|
| | | Total low-coverage genes w/o hypotheticals | Strain-specific genes* | Genes with other copies in genome |
| Lucky Strike | 1534-1685 (45-50%) | 199 (5.8%) | 101 (3.0%) | 98 (2.8%) |
| Semenov | 835-918 (33-36%) | 60 (2.4%) | 30 (1.2%) | 30 (1.2%) |
| Clueless | 929-1111 (30-36%) | 132 (4.3%) | 58 (1.9%) | 74 (2.4%) |
| Lilliput | 1442-1553 (42-45%) | 122 (3.6%) | 60 (1.8%) | 62 (1.8%) |

* detailed information in Tab S.6

**Extended Data Table 2 | Comparison of reported SNP densities from published studies to our present dataset.** Data from *Bathymodiolus* SOX symbionts of this study are depicted in bold. Many of reported values were previously summed up in Wilmes *et al.* (2009). *NA* = information was missing or could not be retrieved.

| Organism | SNPs/kbp | Tool | Environment | Technology | Coverage | Reference |
|---|---|---|---|---|---|---|
| *Ca*. Accumulibacter phosphatis | 0.01 (US)<br>0.02 (OZ) | Consed | Sludge bioreactor | *NA* | 9.2-17.5 x (US)<br>5.36-7.68 x (OZ) | (Kunin et al., 2008) |
| Leptospirillum group II type UBA | 0.04 | *NA* | Acid mine drainage | Sanger | 25 x | (Lo et al., 2007) |
| *Kuenenia stuttgartiensis* | 0.07 | *NA* | Anammox bioreactor | Sanger | 22 x | (Strous et al., 2006) |
| *Solemya velum* endosymbionts | 0.1-1[*] | Custom, GATK | Shallow water clam | Illumina | 52-1115 x | (Russell and Cavanaugh, 2017) |
| Endoriftia symbionts in *Ridgeia piscesae* | 0.3[*]<br>0.9[*] | VarScan<br>GATK | Hydrothermal vent tubeworm | Illumina | 175 x | (Perez and Juniper, 2017) |
| Leptospirillum group II type 5-way CG | 0.9 | Consed, Strainer | Acid mine drainage | Sanger | 20 x | (Simmons, 2008) |
| *Buchnera aphidicola* endosymbiont | 1.8[*§] | Consed | Insects | Sanger | 9 x (?) | (Ham et al., 2003) |
| ‘Iplasma’ | 2.7 | *NA* | Acid mine drainage | *NA* | 20 x | (Wilmes et al., 2009) |
| Endoriftia symbionts in *Riftia pachyptila* | 2.9[#] | Custom | Hydrothermal vent tubeworm | Sanger | 18.6 x | (Robidart et al., 2008) |
| ‘Eplasma’[&] | 5.3 | *NA* | Acid mine drainage | *NA* | 10 x | (Wilmes et al., 2009) |
| ***Bathymodiolus spp.* SOX symbiont** | **5 - 11[*]** | **GATK** | **Hydrothermal vent mussel** | **Illumina** | **100 x** | **This study** |
| *Prochlorococcus* subpopulations | 12 | Custom | Free-living, marine | SAG, Illumina | *NA* | (Kashtan et al., 2014) |
| Gut microbiome | 7-18[*°] | Custom | Human gut | Illumina | 10-32 400 x | (Schloissnig et al., 2013) |
| *Ferroplasma* type II | 22 | Blastn | Acid mine drainage | Sanger | 10 x | (Tyson et al., 2004) |
| *Ferroplasma* type I | 30 | Blastn | Acid mine drainage | *NA* | 4.5 x | (Allen et al., 2007) |
| Archaeal virus contig from metagenome | 70 | *NA* | Yellowstone hot springs | Sanger | 11 x | (Schoenfeld et al., 2008) |
| Archaeal virus AMDV2 | 270 | Consed, Strainer | Acid mine drainage | *NA* | 17.5 x | (Andersson and Banfield, 2008; Wilmes et al., 2009) |

[*] in single host individuals
[#] gene subset
[§] pooled host individuals
[&] partial assembly
[°] total polymorphism percentage of all sampled species per microbiome

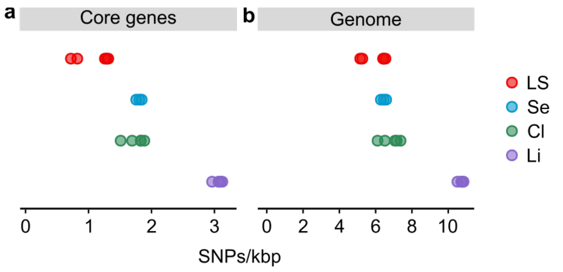


**Extended Data Figure 1 | Single nucleotide polymorphisms (SNPs) of within-host symbiont populations in (a) core genes and (b) whole genome including non-coding regions.** Vent fields (host species) are LS: Lucky Strike (*B. azoricus*), Se: Semenov (*B. puteoserpentis*), Cl: Clueless (*B.* sp.), Li: Lilliput (*B.* sp.).

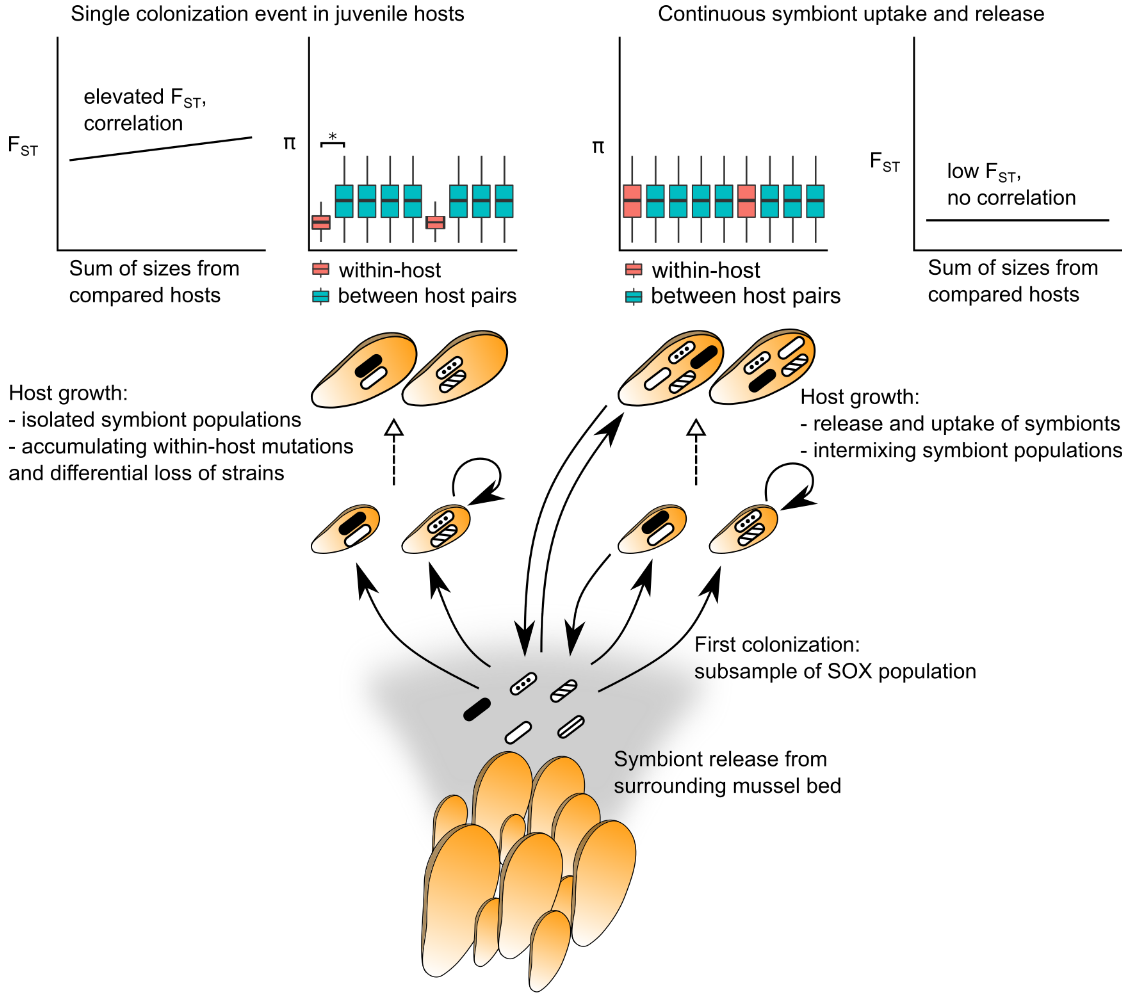


**Extended Data Figure 2 | Theoretical model predicting the influence of symbiont transmission on population genomic signatures (π, $F_{ST}$).** On the left a scenario is depicted in which the symbionts are acquired only once by juvenile mussels during a restricted time window, followed exclusively by repeated self-infection. On the right a scenario is depicted in which the symbionts are continuously released and taken up by host individuals throughout their lifetime.

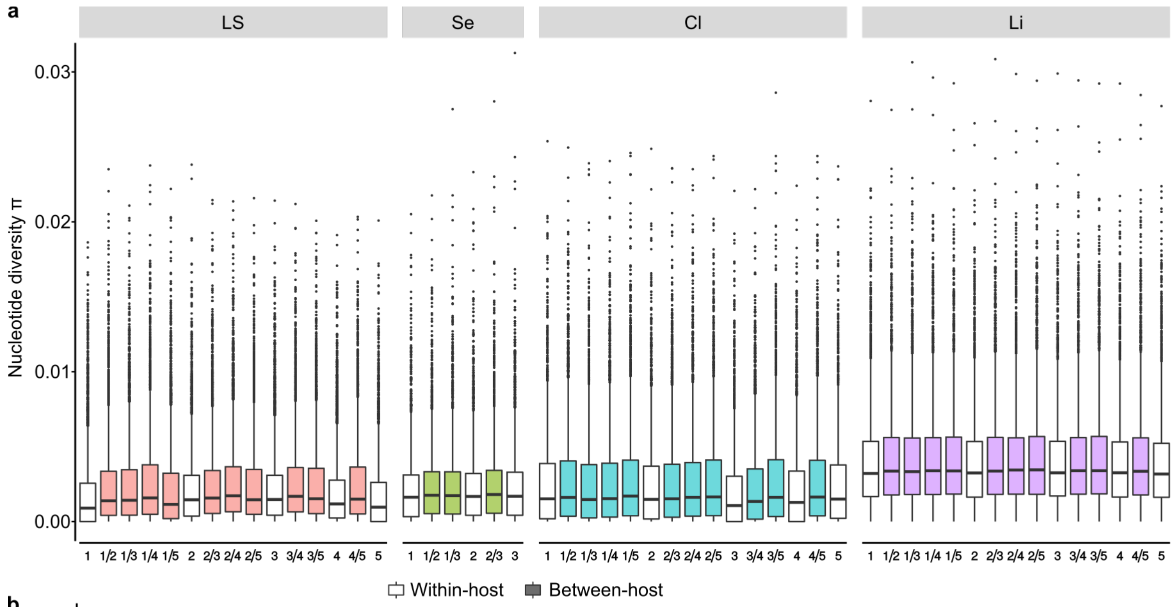


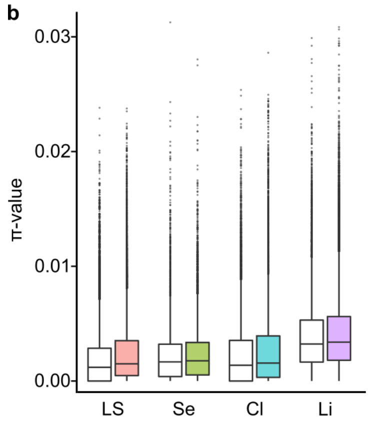


**Extended Data Figure 3 | π-values within symbiont populations of single host individuals (within-host, white) and in pairwise calculation between each two host individuals (pairwise between-host, colored) from all four vent fields.** (**a**) Each within-host π-value per individual and between-host π-value per pair of individuals is shown separately. (**b**) Within-host and between-host π-values are grouped together, respectively. LS: Lucky Strike, Se: Semenov, Cl: Clueless, Li: Lilliput vent fields. In the box-plots, line represents mean, upper and lower hinges represent the first (25th persentile) and third (75th percentile) quartiles, whiskers represent 1.5x interquartile range, individual points are outliers.

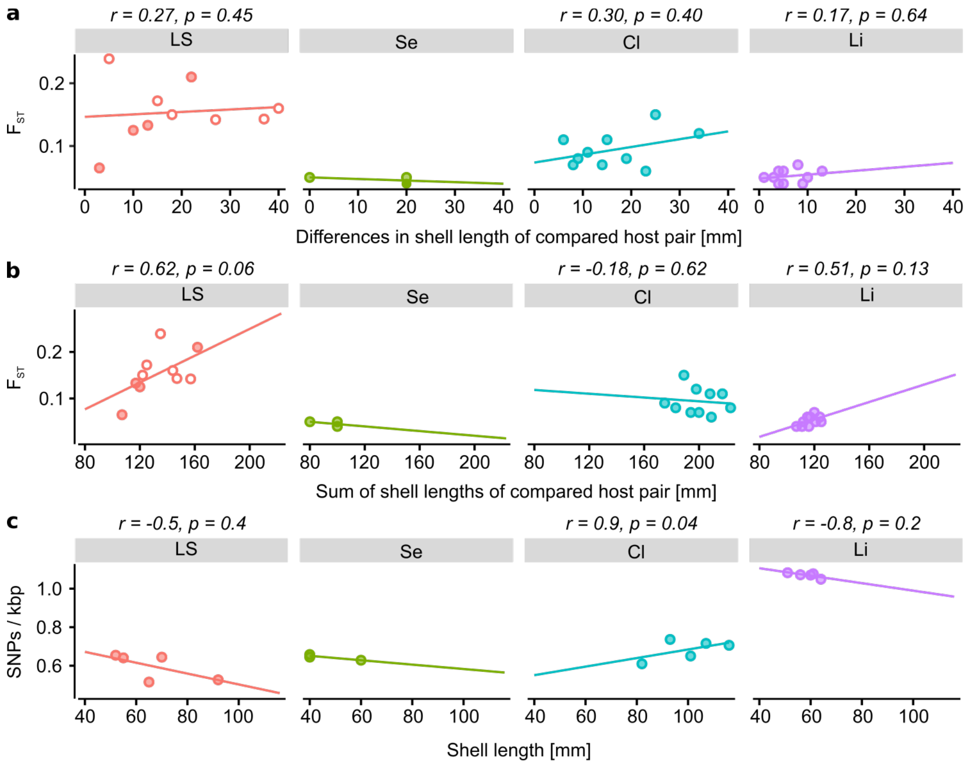


**Extended Data Figure 4 | Spearman correlation between the difference (a) or sum (b) in shell lengths of two compared hosts with the pairwise $F_{ST}$; and correlation of shell length with intra-host SNP density (c).** r = spearmans correlation coefficient rho, p = p-value; for vent field Lucky Strike white symbols = host pairs from different sites Eiffeltower and Montsegur, red symbols = host pairs from the same site; LS: Lucky Strike, Se: Semenov, Cl: Clueless, Li: Lilliput.

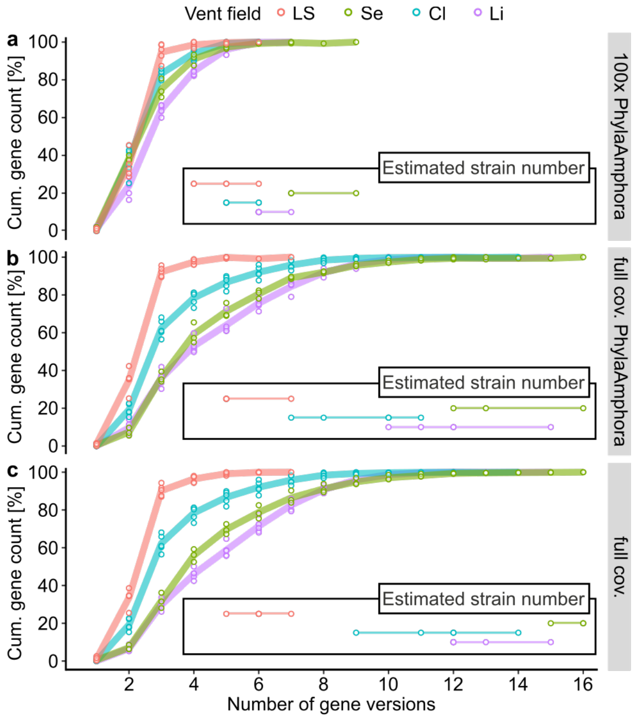


**Extended Data Figure 5 | Cumulative gene counts of distinct numbers of gene versions on gammaproteobacterial marker genes from PhylaAmphora and the extended set of genes that had a read coverage within the coverage range of gammaproteobacterial marker genes, indicating that each strain in the population encoded these.** Strain numbers were estimated for the marker gene set with 100x read coverage (**a**) and full read coverage (**b**) and for the entire gene set with full read coverage per host individual. Full read coverage was 100–120x for LS, 280–370x for Se, 150–215x for Cl and 190–218x for Li mussels. LS: Lucky Strike, Se: Semenov, Cl: Clueless, Li: Lilliput.

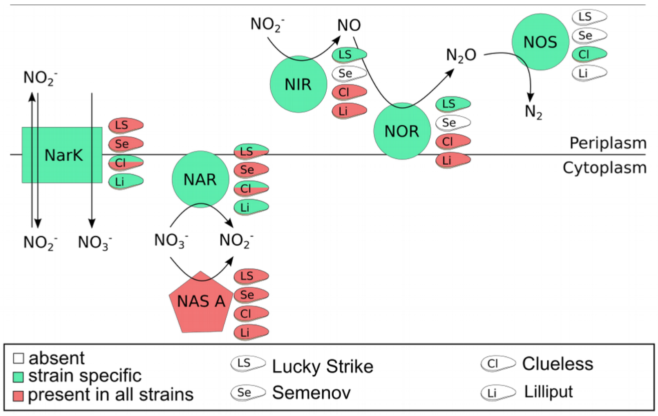


**Extended Data Figure 6 | Representation of denitrification genes among strains of the SOX symbiont.** A gene is absent (white mussel symbols), strain specific (green mussel symbols), or present in all strains (red mussel symbols) in a single host individual. When some host individuals from the same vent site had symbiont populations where a gene is strain specific and others where the entire population encoded that gene, mussel symbol is split into red and green color. LS: Lucky Strike (*B. azoricus*), Se: Semenov (*B. puteoserpentis*), Cl: Clueless (*B.* sp.), Li: Lilliput (*B.* sp.), NAR: respiratory nitrate reductase, NIR: nitrite reductase, NOR: nitric oxide reductase, NOS: nitrous oxide reductase, NarK: nitrate transporter, NAS A: assimilatory nitrate reductase.

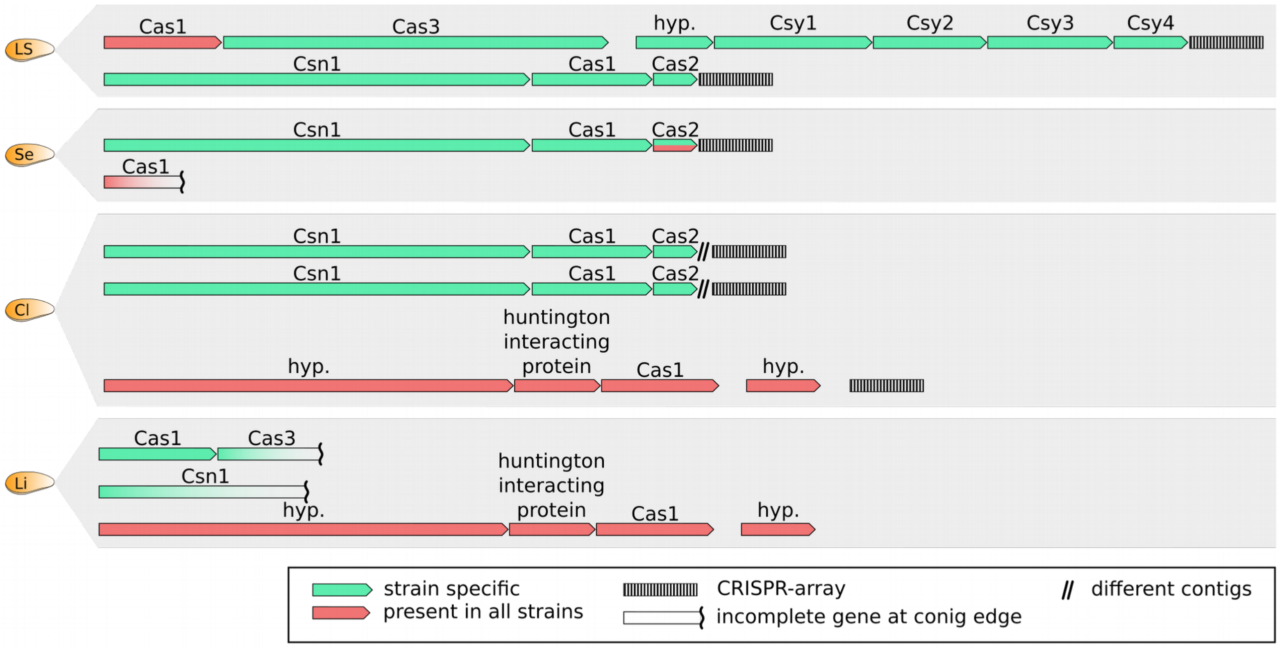


**Extended Data Figure 7 | Representation of CRISPR-Cas gene clusters in the SOX symbiont strains showing strain-specific genes (green), genes present in all strains (red), and CRISPR-arrays (striped boxes).** LS: Lucky Strike, Se: Semenov, Cl: Clueless, Li: Lilliput.
