## Supplementary Material for "Diversity matters: Deep-sea mussels harbor multiple symbiont strains"

### Supplementary Information

#### 1. Methods

**Table S7 | Sample overview.**

| Host species | Vent fields | Mussel sizes [mm] | Year (Cruise) | Depth [m] |
|---|---|---|---|---|
| *B. azoricus* | Lucky Strike (2 sites) | | 2013 (BioBaz) | |
| | Montsegur (MS, 3 mussels) | 65, 55, 52 | | ET 1690 |
| | Eiffeltower (ET, 2 mussels) | 70, 92 | | MS 1700 |
| *B. puteoserpentis* | Semenov-2 | 60, 40, 40 | 2014 (ODEMAR) | 2432 |
| | Ash lighthouse | | | |
| *B.* sp. | Clueless | 93, 116, 82,101, 107 | 2009 (M78-2) | 2972 |
| *B.* sp. | Lilliput (2 dives) | 60, 56, 51, 61, 64 | 2009 (M78-2) | 1490 |
| *B.* sp. | Wideawake | - | 2008 (ATA57) | 2989 |

##### *1.1 DNA and RNA extraction*

Nucleic acids (DNA and RNA) were extracted from the same gill of *B. azoricus* from Lucky Strike with the AllPrep DNA/RNAMini Kit (Qiagen, Germany) according to manufacturer's instructions[1]. Cell debris was removed with QIAshreder Mini Spin Columns (Qiagen, Hilden, Germany). RNA was transcribed into cDNA, using the Ovation RNA-Seq System V2 (NuGEN, USA) and libraries generated with DNA library prep kit for Illumina (Biolabs, Germany). From Semenov, Clueless and Lilliput samples DNA was extracted with the Blood and Tissue Kit (Qiagen, Germany). RNA from Semenov, Clueless and Lilliput samples was extracted from a separate RNAlater-fixed gill tissue sample deriving from the same gill used for DNA extraction with the AllPrep DNA/RNAMini Kit according to manufacturer's instructions (Qiagen, Germany) (Tab. S8, S9). Symbionts from *B.* sp (Wideawake) gill tissue were enriched by homogenization of the entire gill, centrifugation at 100 xg for 10 min, and serial filtration through 12µm, 5µm, 2µm filters[1]. DNA was extracted according to Zhou *et al.*[2].

**Table S8 | Metagenomic sequencing details.**

| DNA | # Individuals | DNA extraction | Seq. Technology | Read length | Sequencing facility |
|---|---|---|---|---|---|
| *B. azoricus* | 5 | AllPrep | HiSeq2500 | 2 x 150bp | MPI Cologne |
| *B. puteoserpentis* | 3 | BloodTissue | MiSeq | 2 x 250bp | MPI Cologne |
| *B.* sp. (Clueless) | 5 | BloodTissue | HiSeq2500 | 2 x 150bp | CeBiTec Bielefeld |
| *B.* sp. (Lilliput) | 5 | BloodTissue | HiSeq2500 | 2 x 150bp | CeBiTec Bielefeld |
| *B.* sp. (Wideawake) | 1 | Zhou et al., (1996) | MiSeq<br>Pacific Biosciences RS II | 2 x 250bp<br>6 SMRT cells | MPI Cologne |

**Table S9 | Metatranscriptomics sequencing details.**

| RNA | # Individuals | RNA extraction | Seq. Technology | Read length | Sequencing facility |
|---|---|---|---|---|---|
| *B. azoricus* | 5 | AllPrep | HiSeq2500 | 2 x 150bp | MPI Cologne |
| *B. puteoserpentis* | 3 | AllPrep | HiSeq2500 | 2 x 150bp | MPI Cologne |
| *B.* sp. (Clueless) | 4 | AllPrep | HiSeq2500 | 2 x 150bp | MPI Cologne |
| *B.* sp. (Lilliput) | 5 | AllPrep | HiSeq2500 | 2 x 150bp | MPI Cologne |

##### *1.2 Sequencing, assembly and annotation*

TruSeq library preparation and sequencing was conducted by the Max Planck Genome Centre in Cologne, Germany, and the Centrum für Biotechnologie (CeBiTec) in Bielefeld, Germany, using Illumina HiSeq2500 with a read length of 150 bp, paired-end, for Lucky Strike, Clueless and Lilliput, and Illumina MiSeq sequencing with read length 250 bp for Semenov and Wideawake (Tab. S8, S9). Before assembly, Illumina TruSeq adapters and the internal Illumina standard for sequencing, phiX, were removed from the raw reads using BBMap (v36.x, Bushnell B. - BBMap – sourceforge.net/projects/bbmap/). Read quality was trimmed to PHRED score 2 and the first 10 bp of all reads were removed.

Metagenomes were assembled according to the following workflow. For each vent field an initial consensus assembly was produced with IDBA-ud (v 1.1.1)[3] with the pooled reads of all mussel individuals of that field. Subsequently, clean reads of each mussel individual were mapped separately to the consensus IDBA assembly to perform differential coverage binning of the SOX symbiont of each mussel individual as implemented in GBtools (v 2.4.5)[4] and the bins further improved with contig connectivity analysis[5]. Clean reads from each mussel individual were then mapped separately to the symbiont bin and reassembled with SPAdes (v 3.1.1)[6] independently to obtain the best bin per host individual. Best individual bins were produced to extract the completest and cleanest read set for the symbiont from each metagenome before a best consensus assembly was produced. When assembly statistics of each individual symbiont bin could not be improved anymore with binning and reassembly, all reads mapping to this optimal bin were pooled and a consensus draft genome per vent field was reassembled using SPAdes[6]. Having a consensus draft genome per field as a reference was essential to be able to compare SNPs across host individuals. The final draft genome statistics, including completeness estimates based on CheckM (v 1.0.7)[7] using gammaproteobacterial 280 marker genes are summarized in Tab. S1. Briefly, symbiont genome sizes were between 2.22 and 2.83 Mb, GC content was around 37.5% and genome completeness was > 94%. Long-read sequencing was done with Pacific Biosciences RS II (PacBio) for the sample from Wideawake (Tab. S7). We sequenced six SMRT cells of extracted DNA from a symbiont enriched fraction. De novo assembly was done using the SMRT software version 2.3.0 with the RS_HGAP_Assembly 2 workflow. We used RAST[8] to annotate the draft genomes we assembled from Illumina and PacBio sequences.

**Table S1 | Details of the SOX symbiont bins.**

| Host species | # Individuals | Completeness* [%] | # Contigs | # CDS | GC [%] | Genome size [Mb] | Read coverage [x] |
|---|---|---|---|---|---|---|---|
| *B. azoricus* | 5 | 94.5 | 607 | 3403 | 37.2 | 2.80 | 101-120 |
| *B. puteoserpentis* | 3 | 94.5 | 250 | 2521 | 37.7 | 2.22 | 281-373 |
| *B.* sp. (Clueless) | 5 | 94.5 | 452 | 3103 | 37.7 | 2.59 | 151-215 |
| *B.* sp. (Lilliput) | 5 | 94.6 | 578 | 3430 | 37.5 | 2.83 | 190-218 |
| B. sp (Wideawake)# | 1 | 94.5 | 379 | 3178 | 37.6 | 2.68 | 162 |

*based on gammabroteobacterial marker genes (Parks *et al.*, 2015)
#from MiSeq symbiont bin

##### *1.3 SNP calling*

For single nucleotide polymorphisms (SNPs) calling clean reads from each sample were further trimmed to a quality PHRED score of 20 and subsequently mapped with minimum identity of 95% to the consensus draft genome from that respective vent field using BBMap. Subsequently, PCR duplicates were removed using samtools (v 1.3.1)[9], reads realigned around INDELs with the Genome Analysis Toolkit (*GATK* v3.3.0)[10] and down-sampled to an average read coverage of 100x for each sample with samtools (per-sample coverages ranged from 100x to 370x).

SNPs were called with GATK HaplotypeCaller, ploidy 10, to allow for polyploidy that accounts for a mixture of multiple coexisting bacterial strains. At ploidy settings above 10 the

SNP numbers did not increase anymore (Tab. S10). Subsequently, unreliable SNPs were filtered with GATK VariantFiltration (settings: QD < 2.0, FS > 60.0, MQ < 40.0, MQRankSum < -20, ReadPosRankSum < -8).

Comparisons of SNP densities with other ploidy settings for all individuals of vent field Clueless showed that lower ploidy settings (2 and 5) reported fewer SNPs whereas a higher ploidy setting of 20 did not increase the detection of SNPs further (Tab. S10). We calculated the SNPs/kbp from the total number of polymorphic sites divided by the consensus genome size per vent field. For calculation of SNPs/kbp in only the core genes, SNPs were extracted for only the ORFs of that gene subset per vent field and divided by the number of nucleotides in the core genes.

To test whether using a per-site consensus reference can introduce artifacts or influence our estimates of symbiont heterogeneity we included an additional control. Using each individuals own symbiont genome bin as reference, we calculated the SNPs/kbp for all individuals from Clueless. All other parameters in the pipeline were kept the same. This revealed that SNP densities with the consensus draft bin as reference made up 93-98% of the single-bin SNP numbers (Tab. S10). This showed that SNP counts were similar, or slightly underestimated, when using the per-site consensus reference. Bash scripts for mapping and SNP calling are provided in https://github.com/rbcan/MARsym_paper.

**Table S10 | SNP densities for the within-host SOX symbiont population of each of 5 host inidivuals from vent field Clueless.** Densities were calculated, using a site-consensus genome assembly (GATK ploidy settings 2, 5, 10 and 20) and each individual-specific symbiont genome bin as reference (GATK ploidy setting 10), respectively to evaluate whether using the consensus bin introduces artifacts. In the present study ploidy setting 10 was used with the consensus reference sequence (marked in grey).

| Host individuals | SNPs / kbp | | | | |
|---|---|---|---|---|---|
| | Consensus reference | | | | Per-individual reference |
| | Ploidy 2 | Ploidy 5 | Ploidy 10 | Ploidy 20 | Ploidy 10 |
| C2 | 6.1 | 7.1 | 7.4 | 7.4 | 7.9 |
| C3 | 5.8 | 6.8 | 7.1 | 7.1 | 7.6 |
| C4 | 4.9 | 5.9 | 6.1 | 6.2 | 6.4 |
| C5 | 5.4 | 6.3 | 6.5 | 6.6 | 6.7 |
| C6 | 5.8 | 6.9 | 7.2 | 7.2 | 7.6 |

*1.4 Core genome calculation and low coverage gene analysis for the detection of strain-specific genes*

The core genome of all SOX symbionts from all four vents, using the consensus draft genome, was estimated with GET_HOMOLOGUES (v1.0)[11], using the OMCL algorithm and a sequence identity setting of 0.5 (Fig. S1). The core genome contained 1283 gene families, representing 40-53% of the predicted ORFs per reference genome.

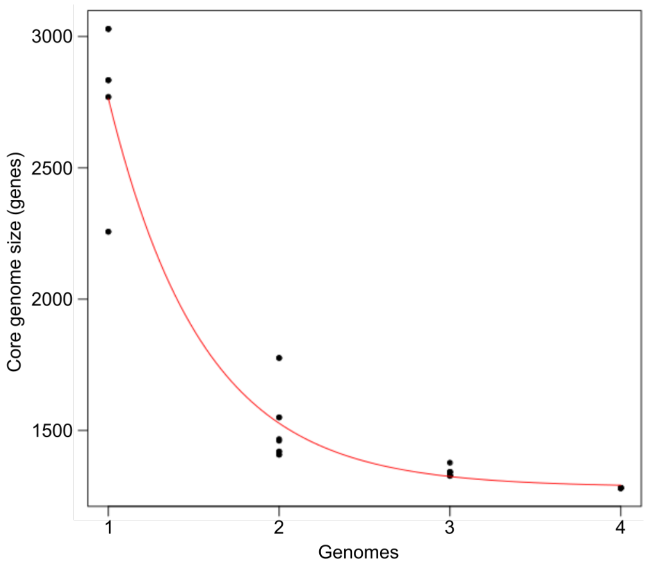


**Figure S1 | Core genome plot of the SOX symbiont along MAR vent fields.** Core genome contains 1283 gene clusters. Gene duplications can lead to different numbers of genes in these gene clusters and we detected 1344 (Lucky Strike), 1350 (Semenov) and 1395(Clueless and Lilliput) genes 1283 clusters. The red line represents the fitted function of the model explaining the decrease in genes that belong to the core genome.

In metagenomes, strain-specific genes not encoded in the genomes of all co-occurring symbionts have a lower coverage compared to single-copy genes which are found in all strains[12]. To systematically identify strain-specific genes, we calculated the per-gene read coverage from our metagenomes. To calculate the per-gene coverage, we used the read mapping files (minimum identity 95%) that were created for the SNP calling (see above). Per-base coverage was calculated with bedtools (v2.16.2)[13], and per-gene coverage for all ORFs was calculated as the mean of the per-base coverages for each ORF with a custom R-script (R version 3.2.2)[14]. To determine the expected coverage of genes that are encoded once by each cell in the population, we extracted more than 200 gammaproteobacterial marker genes (from PhylaAmphora[15]) from the symbiont bins and calculated the coverage for each gene. We plotted the coverage distribution of each gene and of the marker gene set in R (Fig. S3). Some genes had coverage values exceeding those of the single-copy markers, likely resembling unresolved paralogs and repeat elements. All genes that were below the lowest coverage of the marker genes were considered ‘low coverage' and potentially strain-specific.

Low-coverage gene sequences that reached into the last 200 bp of contig edges were not considered because low coverage, as well as high coverage in case of repeat elements, is typical for contig ends and may not reflect strain-specific differences. For functional analysis we excluded all genes that were annotated as “hypothetical proteins”.

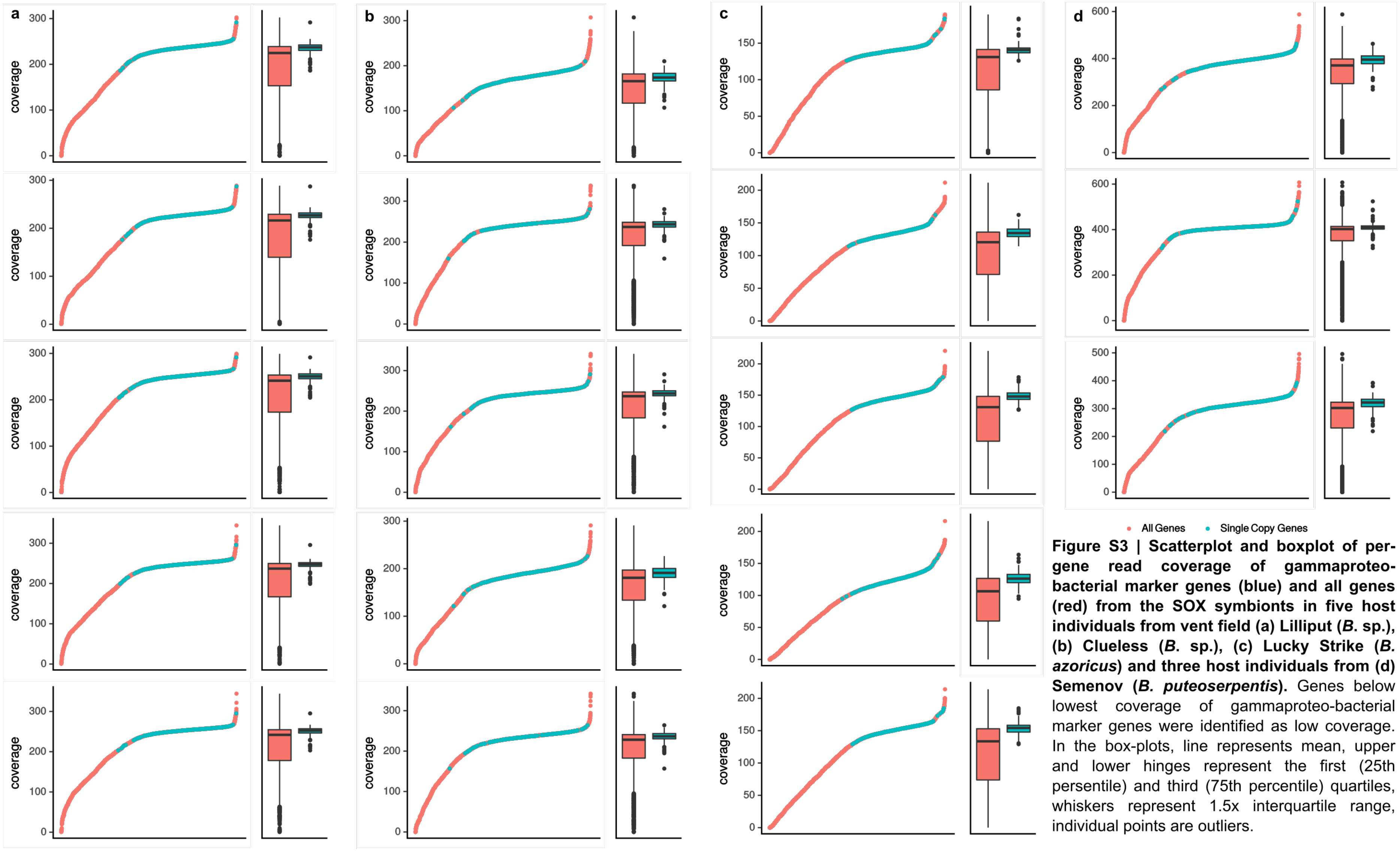


**Figure S3 | Scatterplot and boxplot of per-gene read coverage of gammaproteo-bacterial marker genes (blue) and all genes (red) from the SOX symbionts in five host individuals from vent field (a) Lilliput (*B.* sp.), (b) Clueless (*B.* sp.), (c) Lucky Strike (*B. azoricus*) and three host individuals from (d) Semenov (*B. puteoserpentis*).** Genes below lowest coverage of gammaproteo-bacterial marker genes were identified as low coverage. In the box-plots, line represents mean, upper and lower hinges represent the first (25th persentile) and third (75th percentile) quartiles, whiskers represent 1.5x interquartile range, individual points are outliers.

*1.5 Strain number estimation and test simulation*

Sequencing read coverage of gammaproteobacterial phylogenetic marker genes, as defined by the PhylaAmphora database, was used to detect genes that are present in each strain of the symbiont population as they fell into the same read coverage range (see Method section *Low coverage gene analysis)*. Gene versions were reconstructed for PhylaAmphora markers, as well as all genes encoded by each strain in the population using the tool ViQuaS (v 1.3)[16]. We applied the recommended tool settings for bacterial genomes and discarded all reconstructed sequences below the relative strain frequency above which the haplotype reconstruction is reliable ($f_{min}$) and which was calculated for each gene (Bash scripts in https://github.com/rbcan/MARsym_paper). The highest number of versions for a single gene served as estimation of strain numbers in that particular sample.

To evaluate this approach, a test dataset was created from 10 *E. coli* genomes which showed a heterogeneity of 1% when pooled. Illumina sequencing reads were simulated with ART (v. 2.5.8)[17] for each of the genomes according to Tab. S5. All parameters were chosen to be similar to the features in the sequencing data used in this study. The reads of all 10 genomes were pooled in two different ways, even abundances and uneven abundances of contributing strains, but we consider uneven abundances to be more similar to natural populations. Both samples were assembled with SPAdes (v 3.1.1)[6], mapped with BBMap at 95% identity and subjected to the strain estimation pipeline as described above. The latter was applied to two read coverage depths, 100x and 300x on average, to reach best comparability to our sample data. The strain number estimation from the PacBio assembly was conducted by counting the number of contigs per gene that had full read coverage (defined as above) and were present in one copy only in the Illumina draft genome annotation from the same sample.

**Table S5 | Test dataset, created from the listed 10 *E. coli* strain genomes.** Reads were simulated with ART[#] with default settings unless specified otherwise. Two conditions of uneven and even strain abundances were created (relative abundances shown in table).

| | Strain | BioSample° | Accession° | Settings in ART[#] |
|---|---|---|---|---|
| *Escherichia coli* | P12b | SAMN02603904 | CP002291.1 | Seq. system: HiSeq2500<br>Error: Illumina / default<br>Read length: 150bp<br>Reads: paired-end<br>Insert size: 215bp<br>Min/max quality: 26/40 |
| | ATCC 8739 | SAMN02598405 | CP000946.1 | |
| | K-12 substr. DH10B | SAMN02604262 | CP000948. | |
| | HS | SAMN02604037 | CP000802.1 | |
| | K-12 substr. MG1655 | SAMN02604091 | U00096.3 | |
| | DH1 | SAMN02598470 | CP001637.1 | |
| | REL606 | SAMN02603421 | CP000819.1 | |
| | K-12 substr. BW2952 | SAMN02603900 | CP001396.1 | |
| | BL21-Gold(DE3)pLysS AG | SAMN00002656 | CP001665.1 | |
| | BL21(DE3) | SAMN02603478 | CP001509.3 | |
| Heterogeneity (pooled) | 1%[*] | | | |
| Uneven strain abundances | 1, 0.5, 0.25, 0.125, 0.063, 0.031, 0.016, 0.008, 0.004, 0.002 | | | |
| Even strain abundances | 0.2 all strains | | | |

[*]calculated with Parsnp v. 1.1.2 (Treangen *et al.*, 2014)
°NCBI https://www.ncbi.nlm.nih.gov/
[#]tool ART (Huang *et al.*, 2012)

##### *1.6 Population structure*

SNP frequencies for each sample were extracted from the vcf-files created by our SNP calling analysis.

The $F_{ST}$ calculation is based on Schloissnig *et al.*[17]. Briefly, nucleotide diversity (π) within single host individuals (referred to as $\pi_{within}$) was calculated per gene according to

$$\pi_{s,g} = \frac{1}{|g|} \sum_{i=1}^{|g|} \sum_{n_1 \in \{A,C,T,G\}} \sum_{n_2 \in \{A,C,T,G\} \setminus n_1} \frac{x_{i,n_1,s}\, x_{i,n_2,s}}{c_{i,s}(c_{i,s}-1)}$$

where $s$ is the sample, $g$ is the gene, $|g|$ is the length of the gene, $c_{i,s}$ is the coverage at position $i$ in sample $s$ and $x_{i,nj,s}$ is the number of nucleotides $n_j$ at position $i$ in sample $s$.

Correspondingly, the pairwise between hosts diversity (referred to as $\pi_{between}$) is calculated as

$$\pi_{s_1,s_2,g} = \frac{1}{|g|}\sum_{i=1}^{|g|}\sum_{n_1\in\{A,C,T,G\}}\sum_{n_2\in\{A,C,T,G\}\setminus n_1}\frac{x_{i,n_1,s_1}\,x_{i,n_2,s_2}}{c_{i,s_1}c_{i,s_2}}$$

Then the fixation index $F_{ST}$ for all samples *S* is defined by the ratio of the average sample diversity and the average between-sample diversity[18]:

$$F_{ST}(S,g) = 1 - \frac{\frac{1}{|S|}\sum_{s\in S}\pi_{s,g}}{\frac{2}{|S|(|S|-1)}\sum_{s_1\in S}\sum_{s_2\in S\setminus s_1}\pi_{s_1,s_2,g}}$$

where |*S*| is the number of samples.

We calculated pairwise per gene $F_{ST}$-values between symbiont populations of each two host individuals per vent field and mean per-gene $F_{ST}$-values of all host individuals per field. We further calculated average $F_{ST}$ over all genes for each field.

PCA plots of per-gene $\pi_{within}$ and $\pi_{between}$, considering all core genes among the vent fields, were created with R (v.3.2.2)[14] using the *prcomp* function in the *stats* (v 3.2.3) package and *autoplot* function in the *ggplot2* (v 3.0.0) package. We considered our dataset as multivariate with per-gene π representing the variables. We tested if significant differences exist between $\pi_{within}$ and $\pi_{between}$ at a single vent field, as well as whether significant differences in $\pi_{within}$ exist among vent fields. Finally, we performed a pairwise comparison of $\pi_{within}$ between fields to examine which of the fields differed significantly. These tests were performed using permutational multivariate analyses of variance (PERMANOVA, a*donis* function of the *vegan*, v 2.5.2, package) on a Bray-Curtis dissimilarity matrix (*vegdist* function of the *vegan,* package) calculated for the π-values. Our null hypothesis was that there is no difference among the compared groups, which we defined as follows: For the comparison between vent fields, the $\pi_{within}$ of all the mussels at a single field are considered as one group resulting in a total of four groups; for the analysis at each vent field we consider all $\pi_{within}$ as one group and the $\pi_{between}$, as the second group, with the single mussels or pairs as replicates.

To understand whether there is a correlation between (a) the difference in host shell length of each compared two host individuals and pairwise $F_{ST}$-values, (b) the sum of host shell lengths of each compared two host individuals and pairwise $F_{ST}$-values and (c) shell length and within-host SNP density, we used Spearman correlation (*rcorr* package). Spearman‘s correlation coefficient rho, referred to as *r*, and p-values are shown in Extended Data Fig. 4.

**Table S2 | Average nucleotide diversity π among individuals of the same vent field.** Standard deviation is indicated in the brackets.

| Vent field | Mean π | Within-host π | Pairwise between-host π |
|---|---|---|---|
| Lucky Strike | 0.0024 (± 0.00022) | 0.0022 (± 0.00022) | 0.0026 (± 0.00012) |
| Semenov | 0.0024 (± 0.00007) | 0.0023 (± 0.00006) | 0.0025 (± 0.00005) |
| Clueless | 0.0026 (± 0.00022) | 0.0024 (± 0.00022) | 0.0027 (± 0.00013) |
| Lilliput | 0.0040 (± 0.00011) | 0.0039 (± 0.00004) | 0.0041 (± 0.00003) |

**Tab. S3 | PERMANOVA results comparing within-host π-values of symbiont populations between vent fields.** Analysis performed on values of core genes; 3 degrees of freedom (df) and 14 residual (total df = 17) for the comparison of all 4 vent fields; 1 df and 6 residual (total df = 7) for comparisons of Semenov samples with the other fields and 8 residual (total df = 9) for comparisons between Lucky Strike, Clueless and Lilliput; Pseudo-F = F-value by permutation; P = P-value by 5000 permutations.

| | Lucky Strike | | Semenov | | Clueless | | All | |
|---|---|---|---|---|---|---|---|---|
| | Pseudo-F | P | Pseudo-F | P | Pseudo-F | P | Pseudo-F | P |
| Semenov | 19.18 | 0.018$^{*}$ | | | | | | |
| Clueless | 30.538 | 0.008$^{**}$ | 55.542 | 0.019$^{*}$ | | | | |
| Lilliput | 38.634 | 0.008$^{**}$ | 109.7 | 0.018$^{**}$ | 72.931 | 0.008$^{**}$ | | |
| | | | | | | | 85.822 | **< 0.001**$^{***}$ |

**Tab. S4 | PERMANOVA results comparing within-host π- values of symbiont populations against pairwise between-host π- values of the populations.** Analysis performed on values of all encoded genes with 3609 for Lucky Strike, 2730 for Semenov, 3304 for Clueless and 3640 for Lilliput as well as for core genes; 1 degree of freedom (df) for all; 4 residual (total df = 5) for Semenov and 13 residual (total df = 14) for Lucky Strike, Clueless and Lilliput; Pseudo-F = F-value by permutation; P = P-value by 5000 permutations for Lucky Strike, Clueless and Lilliput and 719 permutations (maximum) for Semenov.

| $\pi_{within}$ / $\pi_{between}$ | All genes | | Core genes | |
|---|---|---|---|---|
| | Pseudo-F | P | Pseudo-F | P |
| Lucky Strike | 1.311 | 0.253 | 0.982 | 0.399 |
| Semenov | 0.321 | 1 | 0.084 | 1 |
| Clueless | 0.848 | 0.529 | 0.721 | 0.602 |
| Lilliput | 0.749 | 0.628 | 0.354 | 0.864 |

#### 2. Results and Discussion

There is some debate about the definition of a microbial ‘strain’ and even species[19]. However, most agree that genotypic differences are required and sufficient for microorganisms to be classified as different strains, without information about phenotypic differences, which can also be found within genetically identical microbial populations[20]. In this study, we define a strain by a distinct sequence version of any coding gene that is encoded by all cells in the symbiont population.

##### *2.1 Population structure of the SOX symbiont*

The population genetic measure $F_{ST}$ can determine how differentiated populations are compared to each other. The $F_{ST}$-values < 0.17 we calculated between symbiot populations from different host individuals deviate substantially from values observed in bacterial populations characterized by clear differentiation. In fact, in the human gut microbiome inter-individual $F_{ST}$ values were 0.4 among most similar and 0.8 among all individuals[21] and in the marine, free-living cyanobacterium *Prochlorococcus* $F_{ST}$ values were ~ 0.7 between subpopulations[22]. Alltogether this data indicate that, despite its intracellular lifestyle, *Bathymodiolus* symbionts exhibit a much higher degree of intermixing than reported for other well studied systems with extracellular or free-living microbial populations of similar heterogeneity levels.

Our extended sampling effort allowed us to investigate data from a range of mussel sizes, which reflect differences in age as shown for other species of bathymodiolin mussels[23]. As shown previously, the gills in *Bathymodiolus* mussels are continuously growing and newly formed gill tissue needs to be colonized by the symbionts throughout the hosts life[24]. However, it was unclear whether this happens exclusively through repeated self-infection within one host or through the uptake of symbionts from the environment, or both. If there is no exchange of symbionts among host individuals, as during repeated self-infection, the symbiont populations of two co-occurring mussels would evolve isolated from each other after the first colonization of juvenile hosts. This will result in a gradual differentiation of the two bacterial populations over time leading to an increase of $F_{ST}$. Accumulation of within-host mutations and differential loss of symbiont strains, decreasing the similarity among populations over time, would result in a positive correlation between the sum of shell lengths of the compared host pair, representing the duration of population isolation, with $F_{ST}$

(Extended Data Fig. 2). We did not detect significant correlations for any vent field, however Lucky Strike samples showed low p-values ($p = 0.06$; Extended Data Fig. 4). Additionally, large and small hosts would have been colonized at very different time points, possibly leading to a correlation between $F_{ST}$ and the difference in host shell lenghts (Extended Data Fig. 1). However, there was no positive correlation of difference in shell length with $F_{ST}$, which could have resulted from two compositionally different seeding populations that colonized the hosts at different times (Extended Data Fig. 4). Instead mussels may continue to take up symbionts from the environment throughout their life. In the case of such continuous symbiont uptake, we do not know whether the symbionts originate from an active free-living population or the release from the surrounding mussel bed. If the symbionts are acquired exclusively from an active free-living population, older individuals would possibly have had contact to other strains that young individuals never encountered. This possibly results in a correlation between heterogeneity and shell length. In the other case where symbionts are acquired from released cells from co-occuring individuals, we would not expect a positive correlation as the symbiont populations would continuously intermix among hosts. We did not detect any significant correlation of heterogeneity with shell length, except for Clueless. The limited number of samples (five) calls for caution when interpreting these results (Extended Data Fig. 4). The fact that we did not detect correlations of $F_{ST}$ with difference in shell length, sum in shell length or between heterogeneity and shell length supports our hypothesis of a continuous uptake mode with intermixing SOX symbiont populations (Extended Data Fig. 4, Extended Data Fig. 2).

Interestingly, at Lucky Strike we observed the least similar symbiont populations among hosts from the same vent field and potentially a correlation between the sum of host shell lengths and $F_{ST}$ values (p-value = 0.06, Extended Data Fig. 4). However, these mussels were sampled from two distinct patches 150 m apart, which might be influencing population dynamics. Typical patchiness of biogeochemical conditions even at small distances at hydrothermal vent fields might favor the higher abundance or presence of certain bacterial strains with advantageous genomic potentials. A recent study by Ho et al.[25], showed high divergence between SOX symbionts of different host individuals, possibly caused by patchiness as the authors detected little divergence when mussels originated from the same patch. This divergence even at a small distance between patches is further support for the continuous exchange of symbionts among mussels that are in close proximity to each other. It also indicates that two different patches might select for different strain compositions.

##### *2.2 Number of strains – test data*

We created a test dataset with simulated reads from 10 *Escherichia coli* strains with 1% nucleotide polymorphisms among them when pooled. We included two scenarios of evenly abundant and differently abundant strains in the population and assembled the consensus genomes for both setups. We assumed that in a natural population, the abundances of different strains are not all even but rather similar to the test with uneven abundances. Following the same gene version construction approach as for the symbiont data we estimated 7 (different abundances) and 10 (even abundances) *E. coli* strains for a coverage of 100x (Fig. S2). Increasing the coverage to 300x, our analysis estimated 12 (different abundances) and 17 (equal abundances) strains.This analysis showed that at a coverage of 300x with even abundances we overestimated the number of strains, but are closest to accurate numbers with uneven strain abundances. For 100x the estimated strain numbers were accurate with even abundances and underestimated with uneven strain abundances.

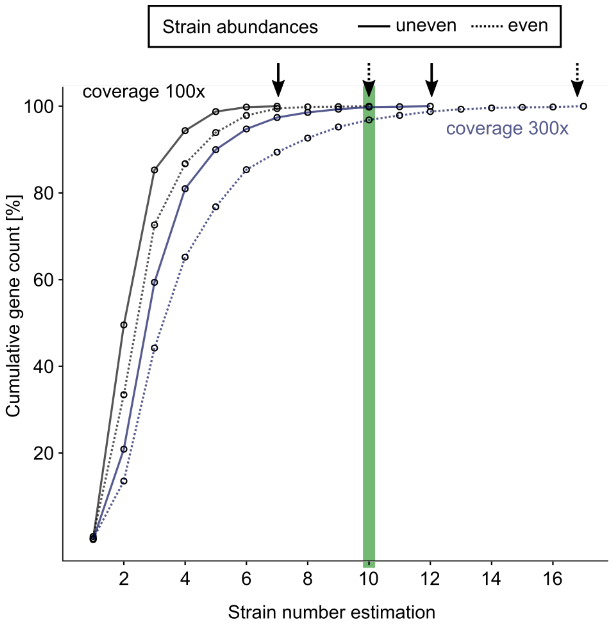


**Figure S2 | Cumulative gene count of numbers of reconstructed gene versions for *E. coli* simulation dataset.** Even (dashed line) and uneven (continuous line) strain abundances are each displayed for 100x (black) and 300x (blue) read coverage, respectively.

*2.3 Accessory genome*

**Table S6 (Excel File) | List of strain-specific genes (all hypothetical proteins were excluded).** List contains gene ID, orthologous family ID, predicted functions, indications of strain-specificity, coverage information [% of single-copy gene coverage], transcript counts [% of single-copy gene mean; TMM normalized]. Purple - Lilliput (*B.* sp. Individuals L102, L104, L105, L51, L54); Green – Clueless (*B.* sp. Individuals C2, C3, C4, C5, C6); Blue – Semenov (*B. puteoserpentis* individuals BputSemA, BputSemB, BputSemC); Red – Lucky Strike (*B. azoricus* individuals BazF, BazG, BazH, BazI, BazJ). In Summary Column: () - other lower covered versions, ‘ - not all lower covered, * - other fully covered version, a – all strains encode it, l – lower covered (strain-specific), x – gene not present; darker color when all (except max. one) individual express it, lighter color when some but not all express it, no shade when none express it (for Lucky Strike samples: all below 10 considered not/low expressed). Expression and gene coverage values are only reported for “l” low-coverage genes. Upper part of the list contains all genes with functional annotation that were “truly” strain-specific at at least one hydrothermal vent field, as no further copy of these genes were identified in the genomes. The lower part of the list reports some additional genes that were detected to have low coverage but had mostly additional low-coverage of full-coverage copies. Here, low coverage might be caused by the fact that different gene neighborhoods (due to genome rearrangements) result in multiple contigs with the same gene in the assembly. For these genes we can not be certain whether every strain carries this gene or not, and considered these genes not “truly” strain specific.

In the following we report details about the distribution of the most prominent categories of strain-specific genes.

**1) Cell-surface**

Multiple genes whose functions were assigned to be involved in the synthesis of cell-surface components, such as O-antigens, and cell-cell interaction mechanisms were strain-specific and expressed at all vent fields. Many of these genes encoded proteins involved in the dTDP-L-Rhamnose biosynthesis pathway. Also, genes involved in biosynthesis of the capsular polysaccharide GDP-L-fucose were strain-specific in Clueless, only lower covered in a few individuals from Lilliput or they had a second low-coverage copy of the same gene at Lucky Strike. These genes encode proteins involved in the biosynthesis of the O-antigen sugar residues. Consistently, previous studies showed that the combination and number of sugar residues in O-antigens are highly variable surface structures and have been reported to be strain-specific[26]. The variability is important as these structures can determine whether a phage can attach to the bacterial cell or whether a cell is being recognized by a particular host. In the SOX symbionts, these genes were often found in clusters and in some cases had no occurrences of SNPs which is further support for their presence in single strains. The second step in O-antigen production is the linkage of the diverse sugars, a process commonly performed by highly specific glycosyl transferases[26]. Consistently, we found glycosyl transferases, possibly involved in these functions, to be strain-specific. Due to their high strain-specificity, O-antigens are commonly used for serotyping to distinguish between medically relevant strains (e.g for *Salmonella* and *Klebsiella*[27–29]). A serotyping approach might provide a good starting point for visualization of the symbiont strain distribution in *Bathymodiolus* gill tissue.

**2) Hydrogen oxidation**

A gene cluster encoding the complete enzymatic machinery for the oxidation of hydrogen, was present at all vent fields. The hydrogenase cluster appeared to be strain-specific at fields Lucky Strike and Clueless with different abundances of the hydrogenase-carrying strain, whereas at the other two fields this cluster was encoded in the entire symbiont population. The membrane-bound group 1 NiFe hydrogenase consists of a large and small subunit of the uptake hydrogenase which are encoded in one gene cluster together with its maturation factors in the SOX symbiont[30,31]. In Clueless mussels, the hydrogenase cluster was encoded by 7-25% of the symbiont population and the small subunit was detected expressed. The other genes in the hydrogenase cluster were barely or not detected in the transcriptome. Conversely, in Lucky Strike symbionts, all genes in the cluster were expressed and based on read coverage we inferred that 40-83% of the symbiont population carried the cluster.

We investigated the link between hydrogen concentrations in the mussel habitats and presence of the hydrogenase gene cluster in symbiont populations. At Semenov, an ultramafic-hosted vent field with hydrogen concentrations in the mM range, most likely all symbiont strains could use hydrogen (unpublished data from Meteor expedition M126 in 2016) and in fact all of them harbored the hydrogenase gene cluster. In contrast, at Clueless, where hydrogen concentrations were below 1 nM[32], only one in ten symbionts had a hydrogenase. Low hydrogen availability, resulting in competition for this limited resource among symbiont strains, would favor the loss of hydrogenase genes, as the cost of maintaining the gene cluster in the genome may outweigh the benefit of additional energy. The link between hydrogen availability and hydrogenase prevalence was not as clear at the other two sampling sites. Lucky Strike had 122 µM hydrogen, and 60% of symbiont strains encoded hydrogenases. Lilliput had 0.87 µM hydrogen, and hydrogenases were detected in all strains. However, hydrogen concentrations alone do not always indicate which energy source is most favorable for free-living microbes, as this may depend on the availability of other energy sources such as sulfide[33]. Additionally, measuring the concentrations of symbiont energy sources such as hydrogen at hydrothermal vents is not trivial, as they can change over time and space even within one mussel bed down to a scale of milimeters[34]. Our data showed that differences in gene content may be tighlty linked to the environment and resource availability, however our ability to predict the precise conditions experienced by the symbiont populations is still influenced by our limitations to accurately determine all environmental parameters.

##### 3) Denitrification

The reduction of nitrate to nitrogen gas ($N_2$) is performed in four steps that require the following enzymes: respiratory nitrate reductase (Nar), nitrite reductase (Nir), nitric oxide reductase (Nor) and nitrous oxide reductase (Nos). We detected a striking variability in the distribution of these genes among coexisting strains (Extended Data Fig. 6). At Lucky Strike, the Nar subunits had variable read coverage representing 60% to 100% of the symbiont population. In contrast, at the same site the genes encoding the enzymes Nir and Nor were strain-specific in all host individuals with a representation of 30 – 75% in the symbiont population. Surprisingly, no strain was capable to perform the last step in nitrate reduction to $N_2$, as the Nos was missing in all strains at Lucky Strike. The entire Semenov symbiont population encoded only the Nar subunits for nitrate reduction to nitrite, but no strain could reduce nitrite to dinitrogen. In Clueless and Lilliput symbionts the Nar was strain-specific (in two out of five host individuals for Clueless and all individuals for Lilliput), while the genes encoding the enzymes Nir and Nor were present in all strains. Clueless was the only vent where the genes for Nos, which performs the last step in nitrate reduction and its two maturation proteins were present. This gene was also strain-specific and encoded in 15 – 50% of the population. Except for the Nos maturation proteins, all genes encoding enzymes involved in the nitrate reduction were expressed at the different vent fields.

It might seem counterintuitive why some strains lack the Nar as it likely provides an advantage during fluctuating oxygen concentrations. However, all strains additionally encode the assimilatory nitrate reductase (NasA) which performs the same reaction and both enzymes can possibly complement each other. A redundancy between NasA and Nar was hypothesized for vesicomyid symbionts where *Ca.* Ruthia magnifica and *Ca.* Vesicomyosocious okutanii both encoded only one of the two enzymes Nar and Nas[35]. Intriguingly, the recently cultivated, free-living relative of the SOX symbiont, *Ca.* T. autotrophicus, was also shown to encode only the Nar and Nor but lacks the enzymes to perform intermediate reactions of nitrate reduction[36]. Unlike the hydrogenase genes, where either the entire gene cluster was present or absent, nitrate reduction genes were much more flexible at the level of single genes in *Bathymodiolus* symbionts. Such a high variation in presence and absence of nitrate reduction genes, in both free-living bacteria and symbionts, suggests that these genes seem very prone to be aquired and lost from genomes. Moreover, this suggests that each of the genes can provide a selective advantage or disadvantage on its own, without necessarily being dependent on the presence of all genes encoding the whole denitrification pathway.

**4) Methanol oxidation**

A gene cluster containing three genes for the oxidation of methanol to formate and four genes for coenzyme pyrroloquinoline (PQQ) synthesis was identified as strain-specific in Lucky Strike symbionts, while present in all strains in Semenov individuals and absent from Clueless and Lilliput symbionts.

**5) $P_i$-dependent gene regulation and $P_i$ uptake**

The high-affinity phosphate transport system PstSCAB and the two-component regulatory system PhoR-PhoB was strain-specific at vent field Lucky Strike and completely absent from vent field Lilliput, whereas it was encoded by all strains at vent fields Semenov and Clueless. Phosphate strongly adsorbs to iron oxyhydroxides. At hydrothermal vents, dissolved phosphate is scavenged by iron minerals released in great quantities by the hydrothermal plume[37,38]. Since iron concentrations differ substantially between vent fields, we would expect that phosphate concentrations in the mussel beds also differ, depending on the iron concentration in the vent plumes. In habitats where phosphate is plentiful, the advantage of a phosphate-dependent regulatory system may not outweigh the cost of maintaining it, thus, it is more likely to be lost. As far as we are aware, phosphate has only been measured at one of our four sampling locations, Lucky Strike. Here, dissolved phosphate concentrations were relatively high when compared to surface seawater (0.6-2.6μM)[39,40]. Hydrothermal fluids at this vent field have relatively low iron concentrations of 60-770 μmol/kg, which is consistent with the inverse correlation existing between iron and phosphate availability[41]. Although no phosphate concentrations have been reported for Lilliput, it is characterized by diffuse venting of low-temperature fluids with low iron concentrations (0.1 – 43 μM)[42]. Considering the interaction between iron and phosphate, this data suggests that less phosphate might be removed by the co-precipitation, resulting in higher phosphate concentrations. The Pho gene cluster provides yet another example of the remarkable flexibility in gene content of the SOX symbiont even among co-existing strains. Moreover, although we cannot draw final conclusions, the examples of hydrogenase and the $P_i$-related genes strongly suggest that the environmental conditions are key drivers of strain diversity and population dynamics.

**6) Phage defense: CRISPR-Cas and RM-systems**

CRISPR-Cas are described as the adaptive immune response of bacteria and archaea towards virus infections. The palindromic repeats are separated by spacers that represent fragments of previously invading virus or plasmid DNA and therefore serve as a memory for a targeted defense during the next infection[43–45]. In Lucky Strike symbionts, two gene clusters contained *cas* genes of the types Cas, Csy and Csn, followed by the CRISPR array of integrated spacer sequences and repeats. All of the *cas* genes were strain-specific with abundances of 30-75%, except for one *cas1* that was present in the entire population (Extended Data Fig. 7). In Semenov symbionts, we identified one CRIPSR-associated gene cluster with Cas1, Cas2 and Csn1 present in 60-91% of the population. No expression of the CRISPR-associated genes could be detected in the transcriptome. At Clueless, two contigs with identical CRISPR-associated genes Cas1, Cas2 and Csn1 were found in 30-90% of the population. These may not be strain-specific but potentially caused by rearrangements around the *cas* genes resulting in different gene neighbourhoods around these genes and thus different contigs during the assembly. The *cas* genes were expressed in some individuals and the Csn1 in all. One additional contig carrying Cas1 was also identified in the entire symbiont population (Extended Data Fig. 7). In Lilliput symbionts no strain specificity could be identified with certainty in the presence of *cas* genes because the genes were detected in contig extremities. Only *cas1* was shown to be present in 41-67% of the population but with a second copy encoded by the entire population (Extended Data Fig. 7).

Further evidence of strain-specific phage pressure is the vast number of restriction-modification (RM) systems we identified in our dataset. RM are phage defense mechanisms, as well, consisting of a restriction enzyme to degrade invading DNA and a methylation unit to protect self-DNA. RM systems of type I, II and III, as well as other genes involved in DNA metabolism appeared in vast numbers in the variable strain genomes with strong indications of genome rearrangements around them (as indicated by mobile elements, transposases and short contig lengths). Furthermore a large fraction of these was expressed, sometimes with variation according to host individual. Besides the function of RM systems in phage defense they were suggested to be involved in other processes such as increase of genomic variation, stabilization of genomic islands, control of HGT among closely related strains and regulation of transcription via methylation causing e.g. altered surface structures[46]. Therefore, all of these may be important factors in the host-symbiont, symbiont-symbiont and symbiont-phage interactions.

*2.4 How does symbiont diversity emerge and how is it maintained?*

Phages are key drivers of strain diversity in many bacterial populations[47]. They can also strongly influence pangenome evolution. There are several examples of horizontal acquisition of ecologically important gene clusters through phage infection, such as photosynthesis genes in *Prochlorococcus*[48]. Recently, sulfur-oxidation genes were discovered in phages that infect free-living SUP05 bacteria, close relatives of the SOX symbiont[49,50]. Our study revealed a number of genes that suggest widespread phage infection in the SOX symbionts. These include surface structure genes, restriction-modification (RM) systems and CRISPR-Cas and all of these were regularly found in the strain-specific pangenome (Extended Data Fig. 7). Strain diversity in surface lipopolysaccharides and *cas* genes possibly represent diverging phage defense mechanisms among symbiont strains in a typical arms race[51]. Most of the strain-specific CRISPR-Cas systems were expressed, which could indicate that despite their intracellular lifestyle, the symbionts still face phage infection. While the mechanisms are yet to be investigated, recent findings showed that phage particles can enter mammalian epithelial cells to infect the intracellular pathogen *Staphylococcus aureus*[52]. Alternatively, CRISPR-Cas can also play a role in host-bacteria interactions through modification of surface structures by targeting self-mRNA for degradation[53,54]. Besides phages, other mechanisms can produce and maintain diversity in the populations, such as genome rearrangements through RM systems and exchange of genetic material via outer membrane vesicles[55,56]. While we have indications for a number of potential processes, the mechanisms of how diversity is maintained in the symbiont populations are yet to be determined and will be important to the understanding of the evolution of these symbioses.

#### References


1. Sayavedra, L. Host-symbiont interactions and metabolism of chemosynthetic symbiosis in deep-sea *Bathymodiolus* mussels. PhD dissertation, *Univ. Bremen* (2016).
2. Zhou, J., Bruns, M. A. & Tiedje, J. M. DNA recovery from soils of diverse composition. *Appl. Environ. Microbiol.* **62,** 316–322 (1996).
3. Peng, Y., Leung, H. C. M., Yiu, S. M. & Chin, F. Y. L. IDBA-UD: a de novo assembler for single-cell and metagenomic sequencing data with highly uneven depth. *Bioinformatics* **28,** 1420–1428 (2012).
4. Seah, B. K. B. & Gruber-Vodicka, H. R. gbtools: Interactive visualization of metagenome bins in R. *Front. Microbiol.* **6,** (2015).
5. Albertsen, M. *et al.* Genome sequences of rare, uncultured bacteria obtained by differential coverage binning of multiple metagenomes. *Nat. Biotechnol.* **31,** 533–538 (2013).
6. Bankevich, A. *et al.* SPAdes: A new genome assembly algorithm and its applications to single-cell sequencing. *J. Comput. Biol.* **19,** 455–477 (2012).
7. Parks, D. H., Imelfort, M., Skennerton, C. T., Hugenholtz, P. & Tyson, G. W. CheckM: assessing the quality of microbial genomes recovered from isolates, single cells, and metagenomes. *Genome Res.* **25,** 1043–1055 (2015).
8. Brettin, T. *et al.* RASTtk: a modular and extensible implementation of the RAST algorithm for building custom annotation pipelines and annotating batches of genomes. *Sci. Rep.* **5,** 8365 (2015).
9. Li, H. *et al.* The Sequence Alignment/Map format and SAMtools. *Bioinformatics* **25,** 2078–2079 (2009).
10. McKenna, A. *et al.* The Genome Analysis Toolkit: a MapReduce framework for analyzing next-generation DNA sequencing data. *Genome Res.* **20,** 1297–1303 (2010).
11. Contreras-Moreira, B. & Vinuesa, P. GET_HOMOLOGUES, a versatile software package for scalable and robust microbial pangenome analysis. *Appl. Environ. Microbiol.* **79,** 7696–7701 (2013).

12. Ikuta, T. *et al.* Heterogeneous composition of key metabolic gene clusters in a vent mussel symbiont population. *ISME J.* **10**, 990-1001 (2015).

13. Quinlan, A. R. & Hall, I. M. BEDTools: a flexible suite of utilities for comparing genomic features. *Bioinformatics* **26,** 841–842 (2010).

14. R Core Team. *A language and environment for statistical computing. R Foundation for Statistical Computing.* (2016).

15. Wang, Z. & Wu, M. A phylum-level bacterial phylogenetic marker database. *Mol. Biol. Evol.* **30,** 1258–1262 (2013).

16. Jayasundara, D. *et al.* ViQuaS: an improved reconstruction pipeline for viral quasispecies spectra generated by next-generation sequencing. *Bioinforma. Oxf. Engl.* **31,** 886–896 (2015).

17. Huang, W., Li, L., Myers, J. R. & Marth, G. T. ART: a next-generation sequencing read simulator. *Bioinformatics* **28,** 593–594 (2012).

18. Hudson, R. R., Slatkin, M. & Maddison, W. P. Estimation of levels of gene flow from DNA sequence data. *Genetics* **132,** 583–589 (1992).

19. Achtman, M. Evolution, population structure, and phylogeography of genetically monomorphic bacterial pathogens. *Annu. Rev. Microbiol.* **62,** 53–70 (2008).

20. Ackermann, M. Microbial individuality in the natural environment. *ISME J.* **7,** 465–467 (2013).

21. Schloissnig, S. *et al.* Genomic variation landscape of the human gut microbiome. *Nature* **493,** 45–50 (2013).

22. Kashtan, N. *et al.* Single-cell genomics reveals hundreds of coexisting subpopulations in wild *Prochlorococcus*. *Science* **344,** 416–420 (2014).

23. Smith, E. B., Scott, K. M., Nix, E. R., Korte, C. & Fisher, C. R. Growth and condition of seep mussels (*Bathymodiolus childressi*) at a Gulf of Mexico brine pool. *Ecology* **81,** 2392–2403 (2000).

24. Wentrup, C., Wendeberg, A., Schimak, M., Borowski, C. & Dubilier, N. Forever competent: deep-sea bivalves are colonized by their chemosynthetic symbionts throughout their lifetime. *Environ. Microbiol.* **16,** 3699–3713 (2014).

25. Ho, P.-T. *et al.* Geographical structure of endosymbiotic bacteria hosted by *Bathymodiolus* mussels at eastern Pacific hydrothermal vents. *BMC Evol. Biol.* **17,** 121 (2017).

26. Samuel, G. & Reeves, P. Biosynthesis of O-antigens: genes and pathways involved in nucleotide sugar precursor synthesis and O-antigen assembly. *Carbohydr. Res.* **338,** 2503–2519 (2003).

27. Blixt, O., Hoffmann, J., Svenson, S. & Norberg, T. Pathogen specific carbohydrate antigen microarrays: a chip for detection of *Salmonella* O-antigen specific antibodies. *Glycoconj. J.* **25,** 27–36 (2008).

28. Erridge, C., Bennett-Guerrero, E. & Poxton, I. R. Structure and function of lipopolysaccharides. *Microbes Infect.* **4,** 837–851 (2002).

29. Hansen, D. S. *et al. Klebsiella pneumoniae* lipopolysaccharide O typing: revision of prototype strains and O-group distribution among clinical isolates from different sources and countries. *J. Clin. Microbiol.* **37,** 56–62 (1999).

30. Petersen, J. M. *et al.* Hydrogen is an energy source for hydrothermal vent symbioses. *Nature* **476,** 176–180 (2011).

31. Vignais, P. M. & Billoud, B. Occurrence, classification, and biological function of hydrogenases: an overview. *Chem. Rev.* **107,** 4206–4272 (2007).

32. Perner, M. *et al.* Short-term microbial and physico-chemical variability in low-temperature hydrothermal fluids near 5°S on the Mid-Atlantic Ridge. *Environ. Microbiol.* **11,** 2526–2541 (2009).

33. Perner, M. *et al.* Linking geology, fluid chemistry, and microbial activity of basalt- and ultramafic-hosted deep-sea hydrothermal vent environments. *Geobiology* **11,** 340–355 (2013).

34. Zielinski, F. U., Gennerich, H.-H., Borowski, C., Wenzhöfer, F. & Dubilier, N. In situ measurements of hydrogen sulfide, oxygen, and temperature in diffuse fluids of an ultramafic-hosted hydrothermal vent field (Logatchev, 14°45′N, Mid-Atlantic Ridge): implications for chemosymbiotic bathymodiolin mussels. *Geochem. Geophys. Geosystems* **12,** Q0AE04 (2011).

35. Kleiner, M., Petersen, J. M. & Dubilier, N. Convergent and divergent evolution of metabolism in sulfur-oxidizing symbionts and the role of horizontal gene transfer. *Curr. Opin. Microbiol.* **15,** 621–631 (2012).

36. Shah, V., Chang, B. X. & Morris, R. M. Cultivation of a chemoautotroph from the SUP05 clade of marine bacteria that produces nitrite and consumes ammonium. *ISME J.* **11,** 263–271 (2017).

37. Feely, R. A., Trefry, J. H., Lebon, G. T. & German, C. R. The relationship between P/Fe and V/Fe ratios in hydrothermal precipitates and dissolved phosphate in seawater. *Geophys. Res. Lett.* **25,** 2253–2256 (1998).

38. Feely, R. A., Trefry, J. H., Massoth, G. J. & Metz, S. A comparison of the scavenging of phosphorus and arsenic from seawater by hydrothermal iron oxyhydroxides in the Atlantic and Pacific Oceans. *Deep Sea Res. Part Oceanogr. Res. Pap.* **38,** 617–623 (1991).

39. Martiny, A. C., Coleman, M. L. & Chisholm, S. W. Phosphate acquisition genes in *Prochlorococcus* ecotypes: evidence for genome-wide adaptation. *Proc. Natl. Acad. Sci. U. S. A.* **103,** 12552–12557 (2006).

40. Sarradin, P.-M., Caprais, J.-C., Riso, R., Kerouel, R. & Aminot, A. Chemical environment of the hydrothermal mussel communities in the Lucky Strike and Menez Gwen vent fields, Mid Atlantic ridge. *Cah. Biol. Mar.* **40,** 93–104 (1999).

41. Von Damm, K. L., Bray, A. M., Buttermore, L. G. & Oosting, S. E. The geochemical controls on vent fluids from the Lucky Strike vent field, Mid-Atlantic Ridge. *Earth Planet. Sci. Lett.* **160,** 521–536 (1998).

42. Haase, K. M. *et al.* Diking, young volcanism and diffuse hydrothermal activity on the southern Mid-Atlantic Ridge: the Lilliput field at 9°33′S. *Mar. Geol.* **266,** 52–64 (2009).

43. Barrangou, R. *et al.* CRISPR provides acquired resistance against viruses in prokaryotes. *Science* **315,** 1709–1712 (2007).

44. Makarova, K. S. *et al.* Evolution and classification of the CRISPR–Cas systems. *Nat. Rev. Microbiol.* **9,** 467–477 (2011).

45. van der Oost, J., Westra, E. R., Jackson, R. N. & Wiedenheft, B. Unravelling the structural and mechanistic basis of CRISPR-Cas systems. *Nat. Rev. Microbiol.* **12,** 479–492 (2014).
46. Vasu, K. & Nagaraja, V. Diverse functions of restriction-modification systems in addition to cellular defense. *Microbiol. Mol. Biol. Rev. MMBR* **77,** 53–72 (2013).
47. Abeles, S. R. *et al.* Microbial diversity in individuals and their household contacts following typical antibiotic courses. *Microbiome* **4,** 39 (2016).
48. Lindell, D. *et al.* Transfer of photosynthesis genes to and from *Prochlorococcus* viruses. *Proc. Natl. Acad. Sci. U. S. A.* **101,** 11013–11018 (2004).
49. Anantharaman, K. *et al.* Sulfur oxidation genes in diverse deep-sea viruses. *Science* **344,** 757–760 (2014).
50. Roux, S. *et al.* Ecology and evolution of viruses infecting uncultivated SUP05 bacteria as revealed by single-cell- and meta-genomics. *eLife* **3,** e03125 (2014).
51. Bertozzi Silva, J., Storms, Z. & Sauvageau, D. Host receptors for bacteriophage adsorption. *FEMS Microbiol. Lett.* **363,** (2016).
52. Zhang, L. *et al.* Intracellular *Staphylococcus aureus* control by virulent bacteriophages within MAC-T bovine mammary epithelial cells. *Antimicrob. Agents Chemother.* **61,** e01990-16 (2017).
53. García-Gutiérrez, E., Almendros, C., Mojica, F. J. M., Guzmán, N. M. & García-Martínez, J. CRISPR content correlates with the pathogenic potential of *Escherichia coli*. *PLOS ONE* **10,** e0131935 (2015).
54. Sampson, T. R. & Weiss, D. S. Alternative roles for CRISPR/Cas systems in bacterial pathogenesis. *PLoS Pathog.* **9,** (2013).
55. Biller, S. J. *et al.* Bacterial vesicles in marine ecosystems. *Science* **343,** 183–186 (2014).
56. Batstone, R. T., Carscadden, K. A., Afkhami, M. E. & Frederickson, M. E. Using niche breadth theory to explain generalization in mutualisms. *Ecology* **99**, 1039–1050 (2018).
